# Antibiotic-adjuvants abolish resistance conferred by the *Staphylococcus aureus* erythromycin resistance methyltransferase in an *Escherichia coli* model

**DOI:** 10.1101/2025.10.01.679839

**Authors:** Taylor S. Barber, John N. Alumasa

## Abstract

Enzyme-mediated resistance is among the main strategies bacteria use to evade antibiotic action. *S*-adenosylmethionine-dependent erythromycin resistance methyltransferases catalyze the methylation of 23S ribosomal RNA in bacteria, causing resistance to macrolides, lincosamides, and streptogramin type-B antibiotics. Given the diversity and number of identified variants of these enzymes, it is vital to devise ways of inhibiting their activity to rescue affected antibiotics. Here, we use computer-aided solvent mapping and virtual screening techniques to identify inhibitors of Erms displaying promising adjuvant properties. We further demonstrate that an *E. coli* model expressing a recombinant *S. aureus* ErmC (*Sa*ErmC) variant causes substantial resistance to representative macrolide and lincosamide antibiotics. Assessment of test compounds using this resistance model revealed candidates displaying promising adjuvant activity when combined with erythromycin or clindamycin. Antibiotic combinations with a principal candidate oxadiazole, JNAL-016, completely blocked *Sa*ErmC-mediated resistance against erythromycin, resulting in an antibiotic-sensitive phenotype in broth microdilution screening assays. This compound also suppressed ErmC activity, allowing erythromycin to regain its bactericidal properties when assessed in actively growing cultures using time-kill assays. JNAL-016 displayed a noncompetitive mode of inhibition against *Sa*ErmC activity in vitro and bound the purified enzyme with high affinity (K_d_ = 1.8 ± 0.7 μM) based on microscale thermophoresis data. Competition experiments suggested that JNAL-016 competes with SAM for its binding pocket on the enzyme, and this compound exhibited no toxicity against human embryonic kidney cells. These findings establish a practical strategy for targeting Erm-mediated resistance, which could lead to a viable adjuvant-based therapy against bacterial pathogens that weaponize variants of this class of methyltransferases.

## INTRODUCTION

The prevalence of infections caused by antibiotic-resistant bacterial pathogens remains a significant health challenge that requires urgent attention, especially given the sharp decline of new antibiotic candidates emerging from the FDA approval pipeline over the past few decades.^1,2^ Current statistics estimate about 2.8 million infections and 35,000 deaths occurring annually from resistant microbes in the U.S., but these numbers are more alarming from a global perspective, with dire projections by 2050 if the problem is left unresolved.^3,4^ Bacterial evolution, including the rampant spread of old and newly acquired resistance mediators, makes tackling this issue daunting. Confronting this task is particularly challenging when considering species with acquired multidrug resistance mechanisms, commonly referred to as superbugs, which often lead to untreatable life-threatening clinical infections.^5,6^ Such microorganisms use myriad strategies to survive antibiotic action, including reduced uptake or efflux of drugs from the cell, and target or antibiotic-modifying enzymes.^7,8^ Given the diversity and complexity of these bacterial defense tactics, it is vital to devise countermeasures for targeting such weaponry to circumvent their survival mechanisms and avoid a looming health catastrophe.

Enzyme-mediated resistance is among the primary strategies pathogenic bacteria use to counteract antibiotic action.^9,10^ These enzymes are mainly located on transmissible mobile genetic elements, like plasmids, which facilitate their rapid inter-species spread through horizontal gene transfer, increasing the health risk level to animals and humans considerably.^10–12^ Resistance-conferring enzymes are typically not essential for bacterial-cell viability, and several classes are currently clinically active, neutralizing the effectiveness of many antibiotics, including those designated as our last line of defense.^13,14^ The nonessential character and prevalence of such enzymes conveniently provide a potential therapeutic strategy for developing antibiotic-adjuvant-based treatment regimens. These treatment options would evade associated resistance mechanisms linked to the affected classes of antibiotics by neutralizing the activity of the enzyme(s) responsible.^15,16^ Ultimately, this approach would restore the activity of doomed antibiotics that succumbed to similar bacterial defenses, allowing precious time for the discovery and development of more efficacious antibiotics to fight resistant pathogens. Also, targeting nonessential pathways would avoid any potential selective pressure, eliminating the requirement for a swift adaptive survival response by bacterial cells. Several prominent, well-documented classes of enzymes facilitate widespread antibiotic resistance, including β-lactamases and aminoglycoside-modifying enzymes.^17–19^ Research over the past few decades on the former class provides a glimpse of the potential effectiveness and application of adjuvant-based options for tackling antibiotic resistance mediated by enzymes.^20,21^

Bacterial methyltransferases are among the known resistance-conferring enzymes. Despite their rapid inter-species spread over the last few decades, their threat level has not gained a deserved high-profile reputation.^22–24^ A subset of these enzymes mainly modify an antibiotic’s molecular target by catalyzing the transfer of a methyl group(s) to corresponding substrates using *S*-adenosylmethionine (SAM) as a cofactor.^25,26^ Most members of this class modify the bacterial ribosomal RNA (rRNA) at strategic positions near the binding sites of various antibiotics, promoting resistance by indirectly inactivating protein synthesis inhibitors.^23^ The resulting modifications arguably cause steric hindrance, which is responsible for the observed resistance phenotype affecting several classes of antibiotics.^26–28^ Erythromycin resistance methyltransferases (Erms) belong to this group of enzymes. They utilize a well-characterized mechanism to mono or di-methylate adenosine 2058 (A_2058_; *E. coli* numbering) of the 23S rRNA in bacterial ribosomes, facilitating resistance to macrolides, lincosamides, and streptogramin type-B antibiotics.^28,29^ Remarkably, the strategic placement of the methylated nucleotides has no adverse impact on ribosome function, which might explain why this type of resistance is on the rise.^30^ Several Erm variants exist in clinical settings,^31^ magnifying the potential threat they pose, especially if they end up in bacterial strains with an already established multidrug resistance phenotype.^32^ ErmC is an example of a well-characterized methyltransferase, which was first discovered in 1971 and has since spread into multiple species of pathogenic bacteria, including *S. aureus* and *Corynebacterium diphtheriae*.^22,33^ ErmC genes found in different species are highly conserved, suggesting that an inhibitor would likely have a broad-spectrum effect against this subclass of enzymes.^34,35^ The limited number of studies investigating strategies for inhibiting this enzyme’s activity highlights the use of computational screening to examine small molecules or peptides as potential inhibitors.^36,37^ Some of these studies further use minimum inhibitory concentration (MIC) data for comparative analysis to validate the biological application of such inhibitors.^36–39^ These studies lack a more comprehensive biological characterization of the identified compounds, highlighting their impact on recovering the activity profiles of the affected antibiotic. For example, there is limited evidence on the ability of inhibitors to revive the bactericidal properties of the antibiotic in question. Such analyses would paint a more vivid picture of how inhibitors of this methyltransferase can be applied clinically to fight this form of resistance. The ever-expanding diversity of Erm variants recorded over the past few decades paints a dire picture of bacterial evolutionary response to antibiotics, with no proper countermeasures in place to tackle this form of resistance. With so many identified variants from this subclass of enzymes, it is essential to devise ways of inhibiting their catalytic activity to rescue affected antibiotics and prevent an imminent health crisis.

Here, we present a comprehensive approach, combining computational and wet-lab methods, to characterize the impact of inhibiting ErmC on the antibacterial activity of affected antibiotics. We employ macromolecular solvent mapping and virtual screening to identify small-molecule binding hotspots and prospective inhibitors of ErmC. We use a constructed *E. coli* resistance working model expressing the *S. aureus* ErmC (*Sa*ErmC) variant to demonstrate substantial loss in activity for erythromycin (ERY) and clindamycin (CLN). Using this strain, we conducted single-dose combinatorial screens and identified test compounds with antibiotic-adjuvant profiles. We further establish that an identified noncompetitive inhibitor of *Sa*ErmC, JNAL-016, binds to the SAM binding pocket and exhibits promising adjuvant properties by terminating ErmC-mediated resistance, restoring the bactericidal properties of ERY. Our findings suggest that targeting resistance-conferring methyltransferases could be a viable strategy for developing adjuvant-based therapy against this resistance mechanism in pathogenic bacteria.

## RESULTS AND DISCUSSION

### Computer-aided solvent mapping and virtual screens identify potential ErmC inhibitors

We initially employed a computational solvent mapping software (FTMap) to identify small-molecule binding hotspots using an available crystal structure of ErmC (**PDB ID: 2ERC**). The mapping algorithm uses standard structurally distinct probes to identify small molecule binding pockets, located in consensus probe cluster sites.^40^ Using the FTMap server, we scanned the entire structure of this enzyme and identified a handful of predicted hotspots, identifiable by the clustering of probe molecules (**Figure 1A & S1**). Not surprisingly, one of the predicted binding hotspots with significant probe clustering represented the SAM binding site (**Figure 1A**), validating the accuracy of the mapping results. Some other key cluster sites were localized at locations associated with rRNA substrate binding,^41^ including the pocket where A_2058_ usually docks for the methylation event (**Figure 1A & S1**). Encouraged by these results, we conducted computer-aided virtual screens to identify chemical scaffolds predicted to bind to the same crystal structure of ErmC. For this study, we focused on an antibacterial screening compound library from the Life Chemicals, Inc. repository, which contains small drug-like molecules designed to potentially have antibacterial properties.

**Figure 1.**
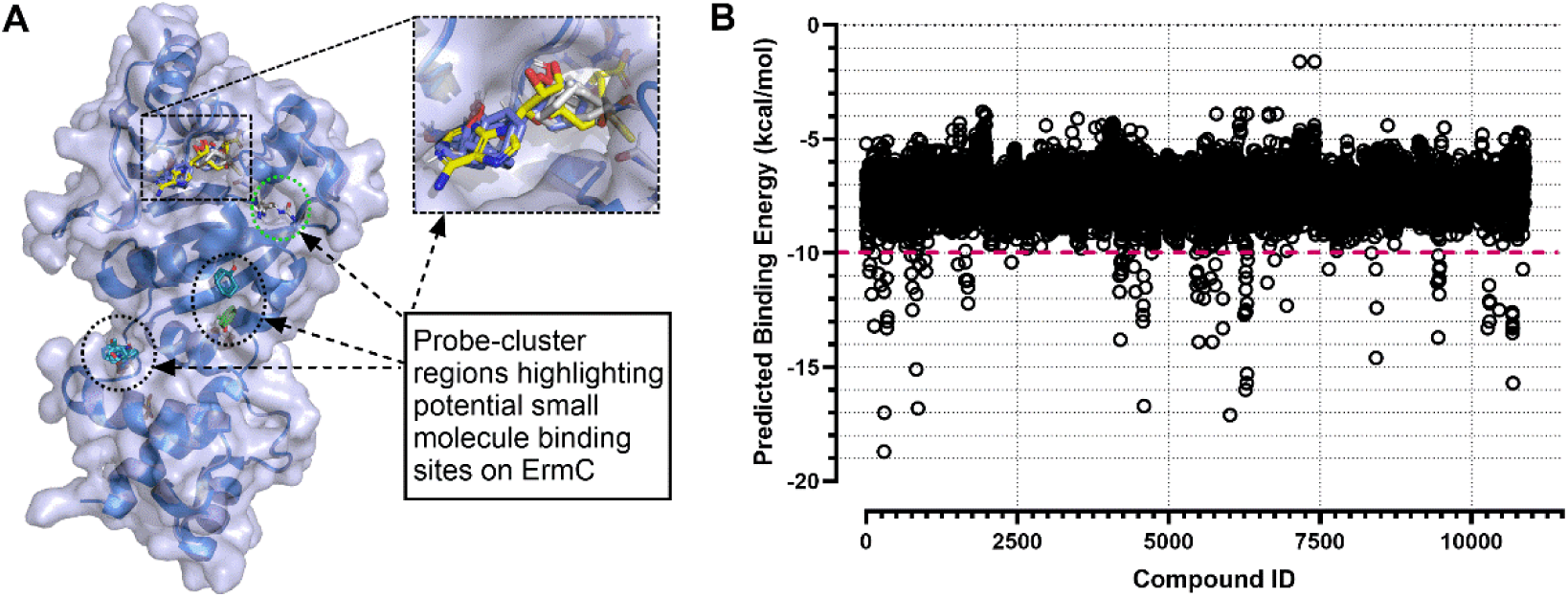
**Solvent mapping and computer-aided screens predict compounds with high binding affinity to *Sa*ErmC**. **A**) Solvent mapping results performed using the FTMap algorithm, highlighting probe cluster regions (dotted sections) representing ‘hotspots’ for potential small molecule binding. The image shows the overlayed structure of an AlphaFold predicted model of the *Sa*ErmC sequence used in this study and the available co-crystal structure of SAM (yellow) bound to the *B. subtilis* ErmC (PDB ID: 1QAO). The green dotted hotspot represents the entry point for A_2058_ to initiate the methylation event. **Inset** – cofactor binding site with SAM highlighted in yellow surrounded by probe clusters, accurately depicting a binding pocket; **B**) Predicted binding energies for identified hits from a virtual screening experiment using AutoDock Vina with the search grid set to scan the entire protein structure. Compounds with a predicted energy ≤-10 kcal/mol were prioritized for secondary computational assessment.

We hypothesized that using such a library would produce inhibitors that can penetrate the bacterial membrane. As part of our strategy, we envisioned using any identified compound exhibiting antibacterial properties at a sub-growth-inhibitory dose for this study. The AutoDock Vina algorithm^42^ was used to conduct an unbiased screen of this library containing 10,880 compounds. The search grid encompassed the entire enzyme structure to increase the possibility of identifying potentially effective inhibitors, based on their binding location. Results from these experiments identified compounds with an array of predicted binding energies, and we focused on those with ≤-10 kcal/mol (**Figure 1B**). This subset of compounds was re-evaluated using the Swiss ADME computational algorithm in a secondary screen for predicting desirable drug-like and pharmacokinetic features.^43^ We compared the compounds based on projected properties, including solubility, lipophilicity, pharmacokinetics, and druglikeness. This assessment enabled us to narrow the candidate list and select a subset of 20 compounds, predicted to bind to the cofactor or substrate binding regions on the protein. We used this subset of compounds for the initial screen, seeking to develop a comprehensive strategy for inhibiting the activity of ErmC and related resistance methyltransferases using antibiotic-adjuvant combinations.

### Expression of *Sa*ErmC in an *E. coli* model causes resistance to macrolides and lincosamides

Variants of ErmC cause resistance against MLS_B_ antibiotics.^44^ To develop a working resistance model for screening potential inhibitors, we codon-optimized the gene sequence for *Sa*ErmC (Uniprot) and cloned it into the tightly regulated arabinose-inducible pBAD24 expression vector. This plasmid was then transformed into an *E. coli* BW25113 strain with a *tolC* efflux pump deletion (BW25113*tolC*::*cat),* resulting in our resistance model strain, TSB-001 (**Table S1**). We confirmed the expression of *Sa*ErmC in this strain following induction with arabinose using SDS PAGE analysis (**Figure S2**). This strain was used to evaluate the resistance phenotype using representative antibiotics ERY and CLN from the macrolide and lincosamide classes, respectively. These antibiotics inhibit protein synthesis by binding to the 50S ribosomal subunit within proximity of A_2058_, hence the observed clinical resistance resulting from methylation of this nucleotide by ErmC variants.^26,44^

Broth dilution assays with these two antibiotics in the presence or absence of *Sa*ErmC expression revealed a dramatic change in their activity profiles. For ERY, expression of this enzyme resulted in 32- and 4.5-fold increase in the recorded MIC and IC_50_ values, respectively (**Figure 2A & B; Table 1**). CLN had a more dramatic change with a recorded 817- and 309-fold increase in its MIC and IC_50_, respectively (**Figure 2C & D; Table 1**), suggesting complete loss of activity under clinical settings. We conducted a control experiment with tetracycline (TET), an antibiotic that inhibits protein synthesis by targeting the accommodation site within the 30S ribosomal subunit.^45^ The binding site of TET is not within proximity of A_2058,_ implying that its activity should be unaffected by the *Sa*ErmC methyltransferase activity. Indeed, the expression of *Sa*ErmC in strain TSB-001 revealed no notable changes in either the MIC or IC_50_ values of TET (**Figure 2E & F; Table 1**). These data suggest that the loss in activity observed for ERY and CLN in the two strains is directly linked to the activity of *Sa*ErmC, which catalyzes the methylation of A_2058_ of rRNA within the 50S bacterial subunit. The data also confirmed the resistance phenotype of the *SaErmC-expressing* strain TSB-001, enabling us to screen for potential adjuvants.

**Figure 2.**
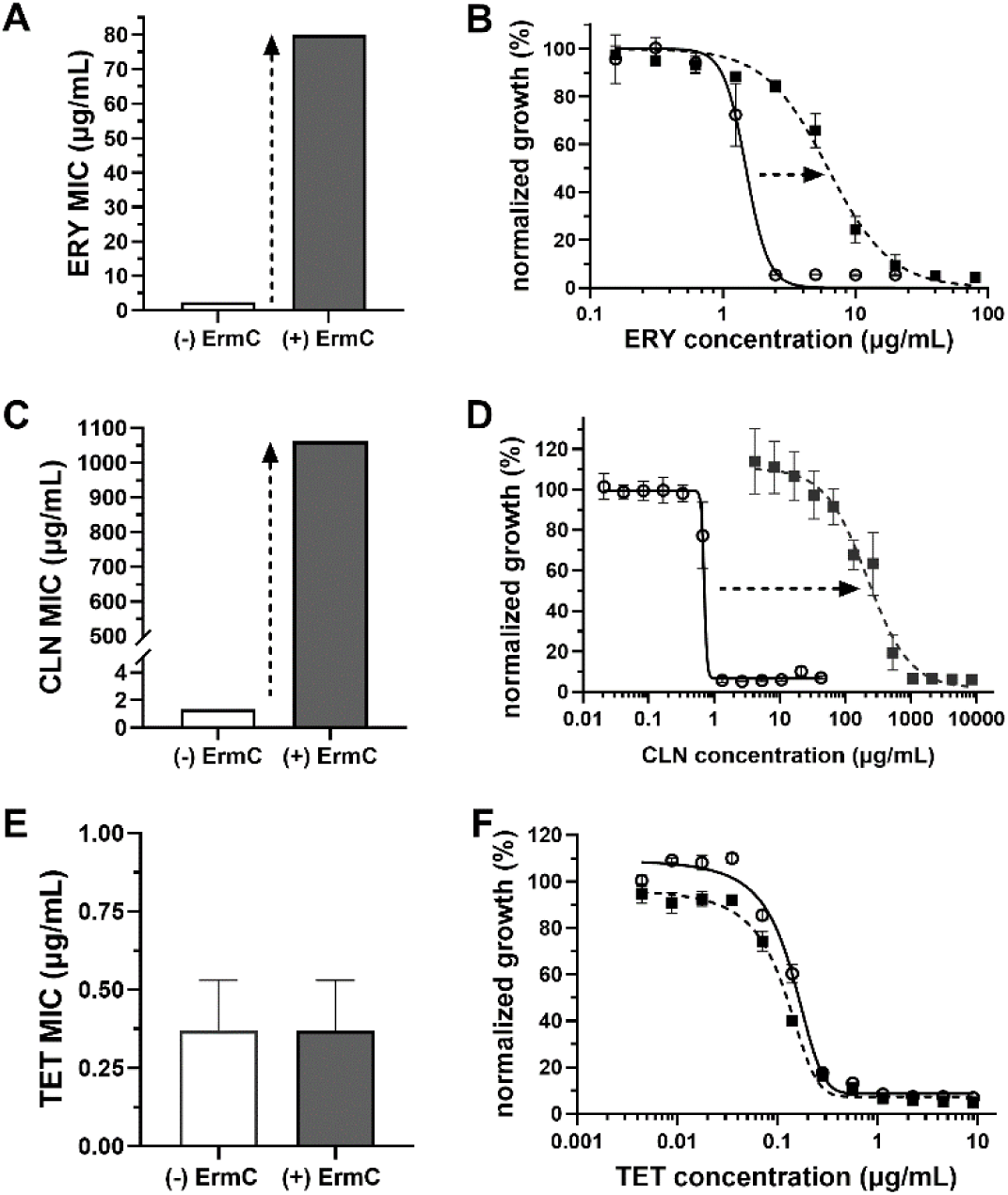
***Sa*ErmC-mediated resistance is specific to MLS_B_ antibiotics**. Results from growth inhibitory experiments conducted with or without *Sa*ErmC expression using broth microdilution assays to determine the minimum growth inhibitory concentrations (**A**, **C**, & **E**) and corresponding dose response profiles (**B**, **D**, & **F**), for ERY (**A** & **B**), CLN (**C** & **D**), and tetracycline (TET) (**E** & **F**) antibiotics. For the dose response plots, the data originates from cells cultured in the presence (filled squares) or absence (open circles) of *Sa*ErmC expression. The dotted arrows indicate a substantial loss in activity of the corresponding antibiotic observed for ERY and CLN but not TET. The data presented are averages from three biological replicates, with the error bars representing the standard deviations.

**Table 1.** Bacterial growth inhibitory activity data for antibiotics against strain TSB-001 in the presence or absence of *Sa*ErmC expression.

| Antibiotic | MIC<sup>a</sup> (μg/ml) |  | MIC fold change | IC<sub>50</sub><sup>a</sup> (μg/ml) |  | IC<sub>50</sub> fold change |
| --- | --- | --- | --- | --- | --- | --- |
|  | (-) SaErmC | (+) SaErmC |  | (-) SaErmC | (+) SaErmC |  |
| CLN | 1.3 ± 0.0 | 1,062 ± 0 | 817 | 0.7 ± 1.7 | 216.3 ± 7.2 | 309 |
| ERY | 2.5 ± 0.0 | 80 ± 0 | 32 | 1.5 ± 2.0 | 6.8 ± 2.4 | 4.5 |
| TET | 0.37 ± 0.16 | 0.37 ± 0.16 | 1.0 | 0.16 ± 0.01 | 1.3 ± 0.01 | 1.3 |

<sup>a</sup>The activity data in the table are presented as the mean ± SD from ≥3 biological replicates

### Cell-based single-dose combinatorial screens identify small molecule candidates with antibiotic-adjuvant properties

We initially set out to evaluate the potential antibacterial properties of the 20 compounds identified from our computational screen, given that they originated from a library of molecules that might possess antibacterial properties. We conducted assays to determine any potential growth inhibitory activity using our *Sa*ErmC *E. coli* resistance model (**Figure 3A**). As presumed, all 20 compounds displayed varying levels of bacterial growth inhibition, some with recorded MIC values below 1 μg/ml (**Table S2**). Given that these small molecules were among those predicted to bind ErmC with high affinity, potentially inhibiting its catalytic activity, we opted to evaluate their potential adjuvant characteristics at much lower concentrations, displaying no considerable growth inhibitory activity for this study. Using our resistance model, we conducted a single-dose combinatorial screen with each compound at 0.02 μg/ml in the presence or absence of ERY (1.25 μg/ml) or CLN (50 μg/ml). Strain TSB-001 was highly resistant to the selected concentrations of both antibiotics, displaying 100% growth under induction conditions (**Figure 2B & D**). Under these conditions, some test compounds (JNAL-9, 11, 20 & 21) still exhibited growth inhibition when tested alone (**Figure 3B**). Regardless, the overall trend for most of the compounds revealed a strong synergistic effect when combined with ERY (**Figure 3C**) or CLN (**Figure 3D**). The combinatorial activity profiles of some test compounds, like JNAL-10, 12, and 16, were particularly encouraging. As a result, we selected six compounds (JNAL-3, 9, 10, 16, 20, 21; **Table S3**) with varying activity profiles and determined an ideal dose for combinatorial assays. When the growth inhibitory profile of ERY was examined in the presence or absence of fixed doses of these prospective adjuvants using our resistance model, all six ERY-inhibitor combinations reduced the MIC and IC_50_ values of the antibiotic by 2 – 5.7-fold, relative to ERY-only values (**Table 2**). The same concentrations of adjuvants decreased the MIC and IC_50_ values of the CLN by 1.4 – 5.7-fold (**Table S4**). These results demonstrate promising antibiotic-adjuvant properties for the test compounds against *Sa*ErmC-mediated resistance. JNAL-016 was particularly attractive given its profile in the initial single-dose screen (**Figure 3B-D**) and its synergistic effect on the MIC of ERY and CLN (**Table 2 & Table S4**, respectively), warranting further characterization.

**Figure 3.**
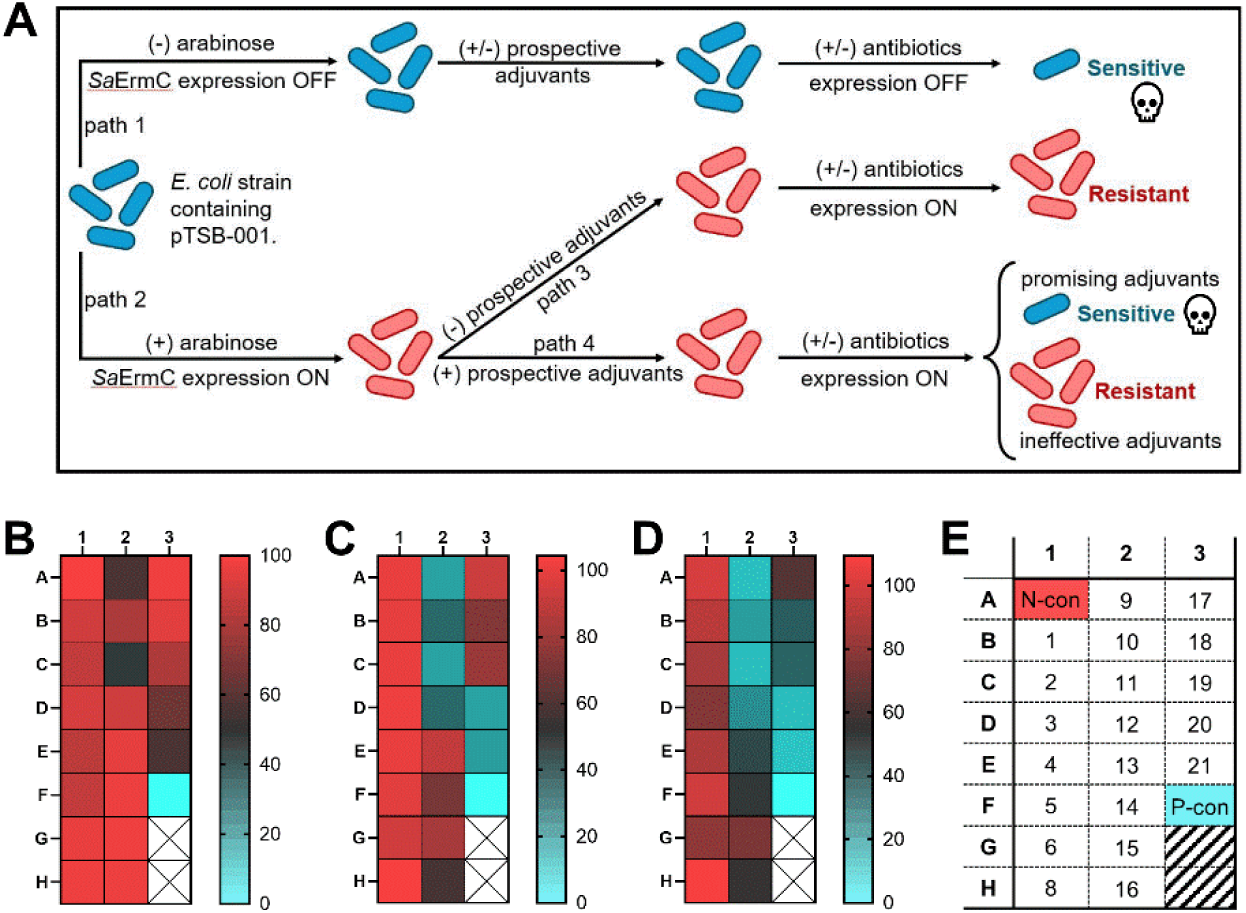
**Cell-based single-dose screen identifies compounds with potential adjuvant properties**. **A**) Schematic showing the design of the cell-based assay used to mimic *Sa*ErmC-mediated resistance for screening prospective antibiotic-adjuvants. Promising adjuvants were selected based on their ability to act synergistically with the corresponding antibiotic and inhibit bacterial growth when expressing *Sa*ErmC. **B-E**) Heat maps showing results from the single dose screening experiments for 20 compounds using the *E. coli* resistance model under *Sa*ErmC expression conditions. The test compounds were each evaluated at a concentration = 0.02 μg/ml. The scales represent the percentage relative growth with the negative control (N-con; red) and the positive control (P-con; cyan) depicting maximum growth and complete inhibition, respectively. The heat-map data highlights the growth inhibitory properties of the test compounds evaluated alone (**B**), or in combination with ERY (1.25 μg/ml) (**C**) or CLN (50 μg/ml) (**D**) at concentrations where the strain exhibits complete resistance. **E**) The identities and location of the test compounds and assay controls in each test plate used to generate the heatmap. Each grid square represents one sample, and the data shown are averages from three-well replicates. Wells G3 and H3 have no samples. See **Table S1** for pTSB-001 description.

**Table 2.** Antibacterial MIC and IC_50_ profiles of ERY in the presence or absence of candidate adjuvants against strain TSB-001 expressing *Sa*ErmC.

| Adjuvant Information |  | ERY MIC (μg/ml) |  |  | ERY IC<sub>50</sub> (μg/ml) |  |  |
| --- | --- | --- | --- | --- | --- | --- | --- |
| ID | Concentration used (μg/ml) | (-) adjuvant | (+) adjuvant | MIC Fold change | (-) adjuvant | (+) adjuvant | IC<sub>50</sub> Fold change |
| JNAL-003 | 0.058 | 80 ± 0 | 40 ± 0 | 2.0 | 6.8 ± 2.4 | 3.1 ± 1.4 | 2.2 |
| JNAL-009 | 0.025 |  | 20 ± 0 | 4.0 |  | 1.6 ± 1.2 | 4.3 |
| JNAL-010 | 0.010 |  | 20 ± 0 | 4.0 |  | 2.9 ± 0.7 | 2.3 |
| JNAL-016 | 0.042 |  | 20 ± 0 | 4.0 |  | 1.2 ± 0.4 | 5.7 |
| JNAL-020 | 0.022 |  | 15 ± 5.8 | 5.3 |  | 2.1 ± 0.7 | 3.2 |
| JNAL-021 | 0.022 |  | 17.5 ± 5 | 4.6 |  | 1.7 ± 0.9 | 4.0 |

<sup>a</sup>The activity data in the table are presented as the mean ± SD from 3 biological replicates

### Prospective adjuvants display no detectable cytotoxicity against human embryonic kidney cells

To assess potential toxicity against mammalian cells, we examined the cidal activity of the six promising adjuvant candidates against human embryonic kidney cells (HEK293) using a propidium iodide (PI) fluorescence permeability cell viability assay.^46^ PI is a charged fluorescent dye impermeable to healthy bacterial membranes but can enter membrane-compromised cells and bind to nucleic acids, enhancing its fluorescence, facilitating cell-death quantification.^46,47^ In this assay, the test compounds were examined by treating cells with concentrations of 20X their corresponding MIC values (**Table S2**). The data obtained was quantified relative to negative control samples, wells containing a cell lysis reagent, which was used to depict 100% death in the assay. The results showed no toxicity for any of the six compounds at the dose examined against HEK293 cells (**Figure S3**). The basal fluorescence detected for all the compound-treated samples was comparable to that of the DMSO-treated samples (≤ 5%). These results provide a promising outlook for developing such compounds for use as adjuvants against *Sa*ErmC-mediated resistance to MLS_B_ antibiotics, because they would provide a large therapeutic window for administering antibiotic-adjuvant combination doses.

### JNAL-016 abolishes the resistance mediated by *Sa*ErmC against ERY

For all follow-up experiments, we used JNAL-016 in this proof-of-concept study. This compound was among those predicted to have the highest binding affinity for *Sa*ErmC from computational data and displayed desirable antibiotic-adjuvant properties in our preliminary cell-based experiments highlighted above (**Figure 3B-D**; **Table 2 & Table S4**). We considered the possibility that JNAL-016 might interfere at the transcription or translation level to prevent expression of the enzyme, indirectly terminating resistance. We quickly disproved this possibility, given that the enzyme was expressed at a comparable level in the presence or absence of this test compound (**Figure S2**). To demonstrate the impact of JNAL-016 on the activity of ERY under *Sa*ErmC-mediated resistance conditions, we conducted a set of growth inhibitory experiments with and without induction of the enzyme in the presence or absence of this compound. The addition of JNAL-016 at 0.042 μg/ml to broth dilution growth inhibitory assays was enough to completely resensitize strain TSB-001 expressing *Sa*ErmC to ERY (**Figure 4A**), terminating the resistance phenotype. However, although the same dose of the adjuvant displayed promising activity when combined with CLN, it did not fully restore sensitivity to this antibiotic (**Figure S4**). This result suggests the need for a higher dose of the adjuvant to overcome the >300-fold increase in the recorded IC_50_ of CLN when the enzyme is expressed (**Figure 2**; **Table 1**). Regardless, these data are promising and illustrate the potential use of adjuvants to tackle *Sa*ErmC-mediated resistance. To further characterize the activity profile of JNAL-016, we complemented this study by conducting growth inhibitory experiments using antibiotic-infused test strips (Liofilchem) on solid agar medium. LB-agar plates were prepared by adding strain TSB-001 in the presence or absence of arabinose into molten top-agar. JNAL-016 (0.25 μg/ml) was added to a subset of plates with *Sa*ErmC expression. We then placed ERY, CLN, or TET-infused test strips onto designated plates to evaluate the potential adjuvant properties of JNA-016 under these growth conditions. Our results confirmed resistance of this strain towards ERY and CLN when *Sa*ErmC was expressed, depicted by a high level of bacterial growth around the test strips at higher antibiotic levels, resulting in elevated MIC values (**Figure 4B & C; Figure S5; Table S5**). The addition of JNAL-016 in the presence of the enzyme reverted the activity of ERY and CLN to mimic those of the antibiotic-sensitive cultures, depicting similar trends to our broth dilution assays (**Figure 4B & C; Figure S5; Table S5**). Importantly, our control experiment demonstrated that the activity of TET was not affected by the presence of *Sa*ErmC or the activity of JNAL-016 (**Figure 4B& C; Figure S5; Table S5**), a result that also confirms the specificity of the Erm-mediated resistance to MLS_B_ antibiotics. We also conducted checkerboard assays to more accurately quantify the synergistic relationship of ERY and CLN when combined with JNAL-016. The cell density data obtained were analyzed using an isobologram and revealed a strong synergistic relationship, whereby 0.03 μg/ml of the adjuvant enhanced the MIC of ERY 16-fold or 4-fold for CLN (**Figure 4D**). This synergistic effect was also evident in the heat maps generated from the cell density data of the checkerboard assays (**Figure S6A & B**). Interestingly, the frequency of spontaneous mutants resistant to a combination of ERY (50 μg/ml) and JNAL-016 (0.042 μg/ml) was <5.1 x 10^-8^, compared to 5.9 x 10^-1^ for ERY alone (50 μg/ml), suggesting that the emergence of resistance to an ERY-JNAL-016 administration would be rare.

**Figure 4.**
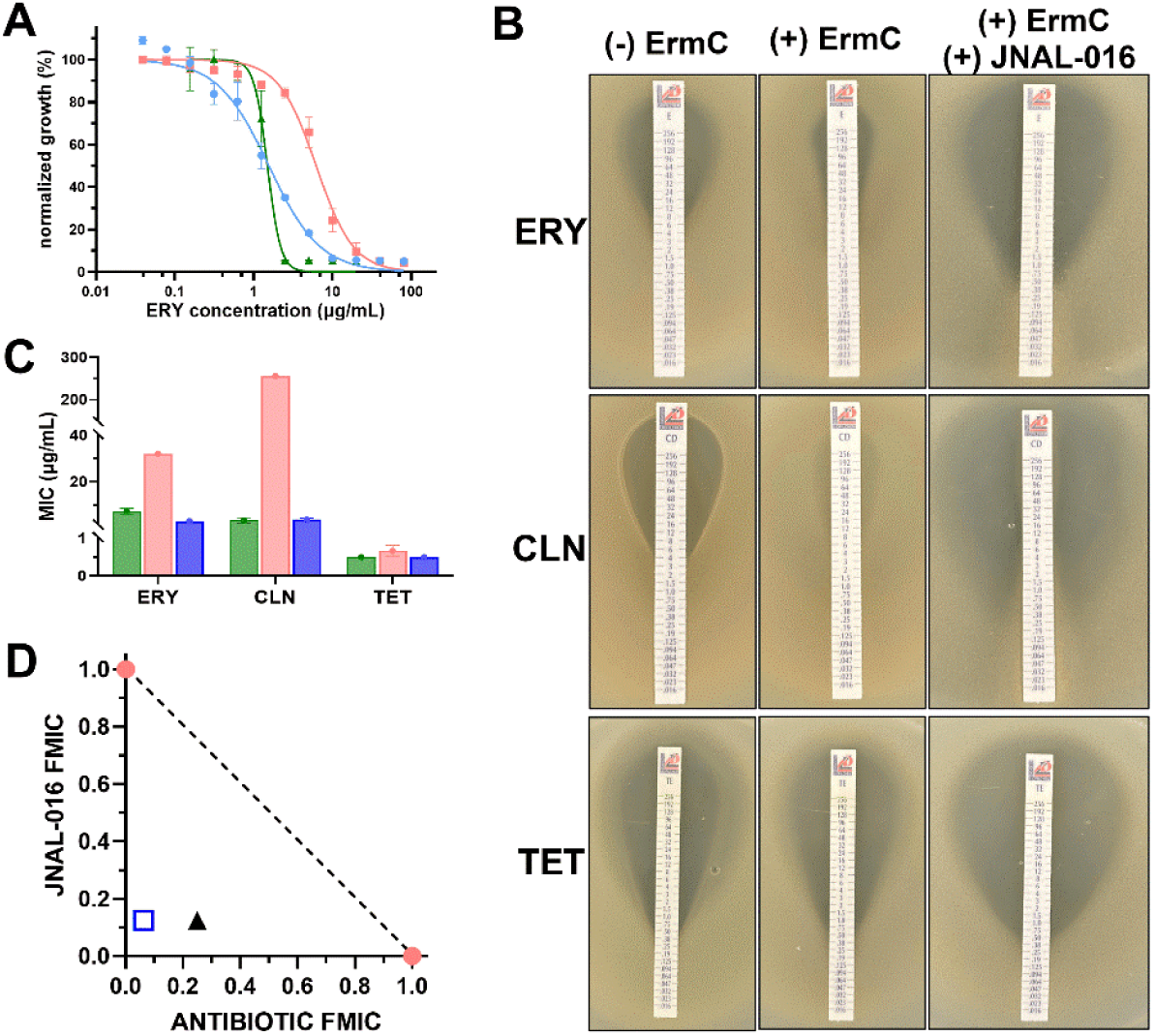
**JNAL-016 displays promising adjuvant properties against *Sa*ErmC-mediated resistance when combined with ERY or CLN**. **A**) Growth inhibitory dose response profiles for ERY assessed in the absence (green triangles) or presence (red squares) of *Sa*ErmC expression and in the presence of both *Sa*ErmC and JNAL-016 (0.042 μg/ml) (blue circles). **B**) Determination of the MICs using antibiotic-infused strips. For induction and samples treated with the adjuvant, arabinose (1%) (middle column) or a combination of arabinose and JNAL-016 (last column) was added to the molten top agar before solidifying it on the plate and placing the strips. The compound concentration did not inhibit growth independently, as evident in the observed bacterial growth at the bottom of the strips. **C**) Graph showing the quantified MIC data from the top-agar experiments. The MICs were determined in the absence (green bars) or presence (red bars) of *Sa*ErmC induction and with an induced set of samples containing JNAL-016 at 0.25 μg/ml (blue bars). For graphs A and C, the data presented are averages from three biological replicates, with the error bars representing the standard deviations. **D**) An isobologram profiling the fractional MIC (FMIC) results from checkerboard assays highlighting the strong synergistic relationship between JNAL-016 and ERY (rectangle) or CLN (triangle) in combination. Only 0.03 μg/ml JNAL-016 was needed to reduce the MIC for ERY by 16-fold (4-fold for CLN). Average data from three biological replicates are presented.

### JNAL-016 displays adjuvant properties against WT BW25113 expressing *Sa*ErmC

We conducted most of our experiments in this study using the efflux-deficient strain TSB-001 (**Table S1**). However, considering that clinical isolates of *E. coli* carrying the ErmC gene would have an active TolC efflux pump, we wanted to establish whether JNAL-016 would retain its adjuvant properties in a wild-type (WT) strain of BW25113, TSB-014, expressing the enzyme (**Table S1**). Initially, we determined the MIC and IC_50_ values for the antibiotics in this strain under induction or repression conditions, and as expected, the values had increased dramatically under both settings (**Table S6**). For example, under ErmC expression, the MIC for ERY against strain TSB-014 increased 100-fold, and that for CLN was 2.5-fold relative to strain TSB-001 (**Table 1 & Table S6**). Checkerboard assays with JNAL-016 revealed a similar synergistic profile for both antibiotics to that observed in the efflux-deficient strain, albeit requiring higher concentrations of the adjuvant (**Figure S6C & D**). For instance, the use of JNAL-016 at a dose 10 times that used to abolish resistance against ERY in strain TSB-001 (**Table 2**), decreased the MIC of this antibiotic 4-fold, or 2-fold for CLN (**Table S6**). These findings are promising when considering clinical isolates of deadly pathogens exhibiting ErmC-mediated resistance. However, follow-up investigations are warranted to fully characterize the activity of this promising adjuvant against clinical isolates.

### Time-kill assays confirm the adjuvant properties of JNAL-016 and its ability to restore the bactericidal activity of ERY

To test the activity of antibiotic-JNAL-016 combinations in actively growing cultures, we employed a PI-based fluorescence detection system in our time-kill assays. For these experiments, PI (5 μM) was added to the culturing medium, and we monitored the synergistic effects of JNAL-016 on the activity of ERY using cultures expressing *Sa*ErmC to mimic the resistance phenotype. Different antibiotic concentrations were evaluated when combined with a fixed dose of JNAL-016 (0.085 μg/ml) in cultures grown at 37 °C in a dry air incubator shaking at 325 rpm for 8 h. During this period, we measured bacterial growth by recording the cell density (OD_600 nm_) and estimated viable cells from the emitted PI fluorescence every hour. To calculate cell death under each concentration of ERY used, we normalized the obtained PI fluorescence by dividing those values by the total cell density to eliminate unreliable data. This strategy accounted for any observed variability in the fluorescence signal resulting from differential growth rates in samples treated with lower vs. higher ERY concentrations. When comparing bacterial growth in the presence or absence of JNAL-016, our experiments demonstrated substantial growth inhibition in samples treated with ERY-JNAL-016 combinations relative to those containing ERY alone (**Figure 5A & B, Figure S7**). This phenomenon was particularly more apparent in samples containing ≤ 20 μg/ml ERY, where ≥ 40% growth (relative to the DMSO control) was recorded in the ERY-only-treated samples (**Figure 5A**). In contrast, the presence of JNAL-016 in samples containing the same range of ERY concentrations showed complete growth inhibition over the 8 h period, except for the later stages of samples containing 0.625 or 1.25 μg/ml ERY, which had elicited some growth (**Figure 5B**). Cell-death profiles had a similar trend, revealing several ERY-JNAL-016 combinations with enhanced cell killing, depicted by an increase in PI fluorescence relative to the DMSO controls (**Figure 5C & D; Figure S7**). The absence of JNAL-016 resulted in varying levels of cell death, proportional to the concentrations of ERY tested (**Figure 5C**). Like the growth profiles, ERY concentrations ≤20 μg/ml caused the least cell death. However, drastic changes were observed in the JNAL-016-treated cultures. While antibiotic-adjuvant combinations involving ERY concentrations ≥40 μg/ml had a similar trend to the ERY-only samples, lower doses of the antibiotic demonstrated the desired adjuvant properties of JNAL-016, enabling ERY to overcome *Sa*ErmC-mediated resistance (**Figure 5D**). Based on the PI cell death quantification, the combined antibiotic-adjuvant effect led to substantial cell death, with ∼90% (1 log unit increase in the fluorescence) reduction in the viable cell count recorded at 2.5 μg/ml ERY after 8 h (**Figure 5D**).

**Figure 5.**
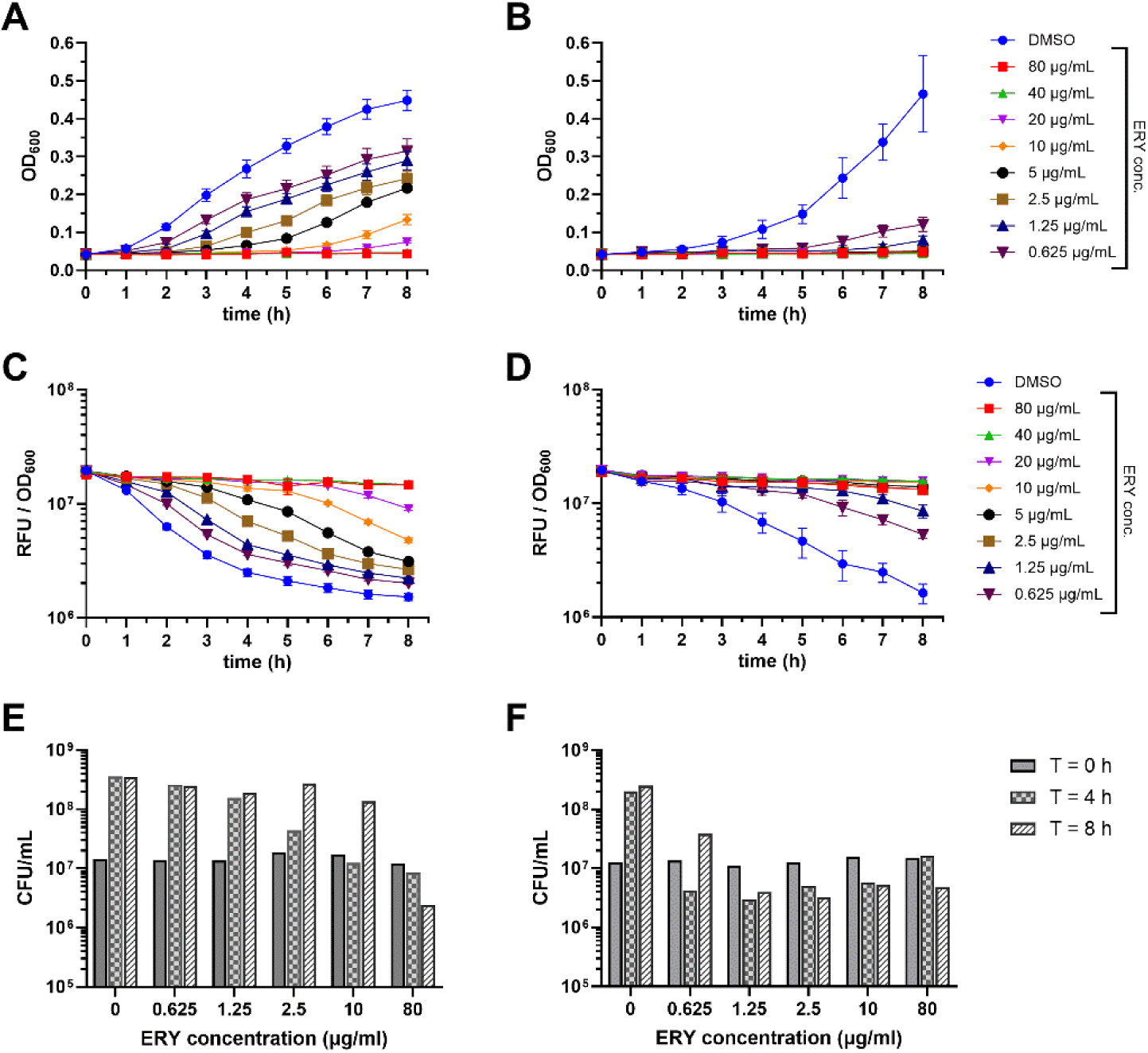
**JNAL-016 acts as an adjuvant to restore the bactericidal properties of ERY in time-kill assays**. **A & B**) Growth profiles of strain TSB-001 expressing *Sa*ErmC treated with varying concentrations of ERY monitored by cell density (OD_600 nm_) every hour over eight hours in the absence (**A**) or presence (**B**) of JNAL-016 (0.085 μg/ml) added to every sample. **C & D**) Time-kill profiles for the same samples in ‘A & B’ presented as cell-density-normalized PI fluorescence over eight hours in the absence (**C**) or presence (**D**) of JNAL-016. Samples with higher fluorescence on the graphs have more dead cells. **E & F**) Quantification of the colony-forming units from the same samples in ‘A & B’ performed at different timepoints and concentrations of ERY in the absence (**E**) or presence (**F**) of JNAL-016. DMSO control cultures for samples in panels A, C & E indicate no ERY present in the sample, while those in panels B, D & F are treated with only JNAL-016 (dissolved in DMSO) at 0.085 μg/ml. For graphs A-D, the data presented are averages from three technical replicates, and the error bars represent the standard deviation of the mean. Additional biological replicates for the time kill assays are presented in **Figure S7**.

To further highlight the restored bactericidal properties of ERY when combined with JNAL-016, we complemented this fluorescence cell viability determination with plating experiments on solid agar to determine viable cell counts. Sample aliquots from ERY or ERY-JNAL-016 combination treatments in the same time-kill assays highlighted above were extracted at 0, 4, and 8 h timepoints, diluted accordingly, and plated on LB-agar. Viable cell counts were quantified after a 24 h incubation period at 37°C. The observed trends from the analyzed samples mimicked those observed in the growth inhibition and PI-linked death profiles (**Figure 5E & F; Figure S7**). When ERY was used alone, 80 μg/ml was the only dose sufficient to kill >99% of the viable cells after 8h. However, this concentration is 32-fold higher than the recorded MIC for this antibiotic in the absence of the enzyme, making it unrealistically high for typical experiments (**Figure 5E**, **Table 1**). At physiologically relevant concentrations, the number of viable cells increased gradually with time, and there was no notable change in the cell counts by the 8 h timepoint for cells treated with ≤10 μg/ml of the antibiotic (**Figure 5E**). In contrast, the addition of JNAL-016 (0.085 μg/ml; a 2x MIC dose was required in active cultures) to the antibiotic-treated cultures led to considerable cell death after 8 h for ERY concentrations ≤10 μg/ml (**Figure 5F**). For example, 2.5 μg/ml of ERY in combination with the adjuvant reduced the number of viable cells by >99% (**Figure 5F**). This concentration is significant because it mimics the recorded MIC for ERY in a strain lacking *Sa*ErmC expression in this study (**Table 1**), confirming the restored bactericidal properties of this antibiotic. Taken together, results from the time-kill assays establish JNAL-016 as a promising candidate for adjuvant-based therapy against *Sa*ErmC-mediated resistance.

### JNAL-016 is a noncompetitive inhibitor of *Sa*ErmC activity in vitro

While JNAL-016 appears to have an alternative antibacterial mode of action at high concentrations, our biological data suggest that when combined with ERY or CLN, it acts as an adjuvant against *Sa*ErmC-mediated resistance at lower doses, lacking independent growth inhibitory activity. To characterize the biochemical interaction and potential inhibitory profile of JNAL-016 against this enzyme, we developed an expression strain for purifying recombinant *Sa*ErmC. The gene encoding *Sa*ErmC was cloned into the pET28a expression vector, which was transformed into BL21(DE3)pLysS cells, resulting in strain TSB-010 (**Table S1**). Expression and purification of *Sa*ErmC from this strain yielded a high-purity protein (**Figure 6A**), which was used to characterize the inhibition kinetics of JNAL-016 using the MTase-Glo methyltransferase assay (Promega). A 32-oligonucleotide (32-nt) RNA fragment from the 23S rRNA, previously shown to be a substrate for different Erm variants,^48,49^ was used to develop an inhibitor screening assay. For this experiment, methylation of the 32-nt RNA substrate catalyzed by *Sa*ErmC using SAM would generate *S*-adenosylhomocysteine (SAH), which is required as a substrate for the reaction that ultimately generates detectable luminescent light.^50^ Initially, we confirmed the linear correlation between the concentration of SAH and the emitted light, which facilitated the quantification of our experimental data (**Figure S8**). The activity of *Sa*ErmC and closely related methyltransferases depends on the universal methyl donor SAM for catalysis. As a result, it is feasible to project that this molecule could have co-purified with our enzyme during purification, potentially interfering with our data analysis. We therefore assessed the purified *Sa*ErmC for possible contamination with co-purified SAM by conducting MTase-Glo activity assays under various conditions, including the absence or presence of exogenously added cofactor (**Figure 6B**). Results from this experiment showed very little catalytic activity (∼5%) in the absence of exogenously added SAM compared to when this cofactor was added to the reaction (**Figure 6B**). We then assessed the steady state kinetics of *Sa*ErmC catalysis and the corresponding inhibition by JNAL-016 using RNA substrate concentrations ranging from 0 to 8 μM over a 30-minute incubation period. *Sa*ErmC catalyzed the methylation of the 32-nt RNA substrate with a V_max_ = 7.6 (±0.42) x 10^-10^ Ms^-1^ and K_m_ = 3.4 (±0.11) x 10^-6^ M (**Figure 6C**, **Table 3**). The addition of JNAL-016 resulted in a 2.1-fold reduction in the reaction rate (*_app_*V_max_ = 3.7 (±0.81) x 10^-10^) but did not alter substrate binding (*_app_*K_m_ = 3.4 (±0.97) x 10^-6^ M) (**Figure 6C**, **Table 3**). Accordingly, this compound lowered the turnover and catalytic efficiency of *Sa*ErmC by ∼2-fold (**Table 3**). A comparison of the kinetic data in the presence or absence of this inhibitor suggested a noncompetitive mode of inhibition. These data demonstrate that JNAL-016 is an effective inhibitor of *Sa*ErmC activity in vitro, supporting the biological data highlighting its adjuvant properties when combined with ERY and CLN against *Sa*ErmC-mediated resistance.

**Figure 6.**
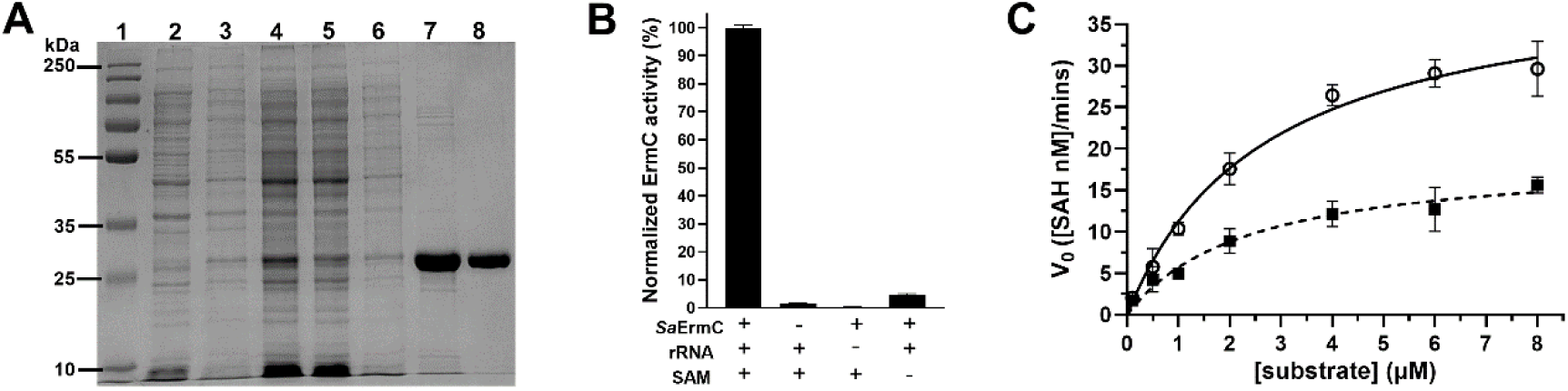
**JNAL-016 is a noncompetitive inhibitor of *Sa*ErmC**. **A**) SDS polyacrylamide gel showing samples from the purification of recombinant *Sa*ErmC following expression in BL21(DE3) cells. L1 – protein marker; L2 – uninduced sample; L3 – induced sample; L4 – clarified lysate; L5 – flowthrough from Ni-NTA binding; L6 – wash fraction after affinity binding; L7 – pooled elution fractions after affinity purification; L8 – purified *Sa*ErmC after gel filtration on an S200 column (Cytiva). **B**) Characterization of the purified enzyme using the MTase-Glo assay to determine the presence of co-purified SAM and potential non-enzymatic auto-hydrolysis of this cofactor under our experimental conditions. The data show no autohydrolysis of SAM in the absence of the RNA substrate, and co-purified SAM accounts for ∼5% of the recovered luminescence signal. **C**) Michaelis-Menten plots showing the steady state kinetics of *Sa*ErmC catalysis for the methylation of a 32-mer RNA substrate in the absence (open circles) or presence (filled squares) of JNAL-016. The data points represent averages from three independent experiments ± SD. These data suggest a noncompetitive mode of inhibition by the inhibitor.

**Table 3.** Michaelis-Menten steady state kinetic data^a^ for *Sa*ErmC in the presence or absence of JNAL-016.

| Sample ID | K<sub>m</sub> (M) | V<sub>max</sub> (Ms<sup>-1</sup>) | K<sub>cat</sub> (s<sup>-1</sup>) | K<sub>cat</sub>/K<sub>m</sub> (M<sup>-1</sup>s<sup>-1</sup>) |
| --- | --- | --- | --- | --- |
| (-) JNAL-016 | 3.4 (0.11) × 10<sup>-6</sup> | 7.6 (0.42) × 10<sup>-10</sup> | 1.5 × 10<sup>-3</sup> | 4.5 × 10<sup>2</sup> |
| (+) JNAL-016 | 3.4 (0.97) × 10<sup>-6</sup> | 3.7 (0.81) × 10<sup>-10</sup> | 7.3 × 10<sup>-4</sup> | 2.2 × 10<sup>2</sup> |

<sup>a</sup>The data presented are the mean values from three independent experiments (SD)

### JNAL-016 competes with SAM, but not the substrate, for the same binding pocket on *Sa*ErmC

The steady state kinetic data for *Sa*ErmC in the presence of JNAL-016 suggested that the compound could bind at an allosteric site to inhibit catalysis. This observation prompted structural studies attempting to co-crystallize an enzyme-compound complex to gain further insight into the binding location and mechanism of inhibition. Although the crystallization attempts have been unsuccessful, we took a different approach that utilized binding experiments to better understand the potential enzyme-inhibitor interactions. We used microscale thermophoresis to conduct a binding experiment titrating JNAL-016 into a fixed enzyme concentration and obtained a binding profile for this compound, which suggested a tight interaction (K_d_ = 1.8 ± 0.7 μM) (**Figure 7A**). To gain further insight into where this inhibitor might be binding, we considered its established noncompetitive mode of inhibition of *Sa*ErmC activity. We then hypothesized that JNAL-016 could either compete with the SAM for its binding pocket or bind at an alternative allosteric site that indirectly blocks cofactor access to the enzyme. To test this hypothesis, we conducted binding competition experiments using microscale thermophoresis, where a fixed concentration (2 μM) of 50-[N-[(3S)-3-Amino-carboxypropyl]-N-methylamino]-50-deoxyadenosine (aza-SAM) was added to serially diluted samples of JNAL-016 prior to adding the enzyme. The labile methyl-sulfonium ion of SAM made it unstable under our binding experimental conditions. As such, we used a more stable analog, aza-SAM, which contains a non-extractable aminomethyl replacement.^51^ This analog has been used in the past to replace SAM in crystallography and other reactions involving methyltransferases.^51,52^ Under these conditions, the enzyme-JNAL-016 interaction decreased substantially, resulting in a ∼57-fold decrease in the binding affinity (*_app_*K_d_ 122 ± 13 μM) (**Figure 7B**). This result suggested a competitive event for the SAM binding site, which reduced the available binding sites for JNAL-016.

**Figure 7.**
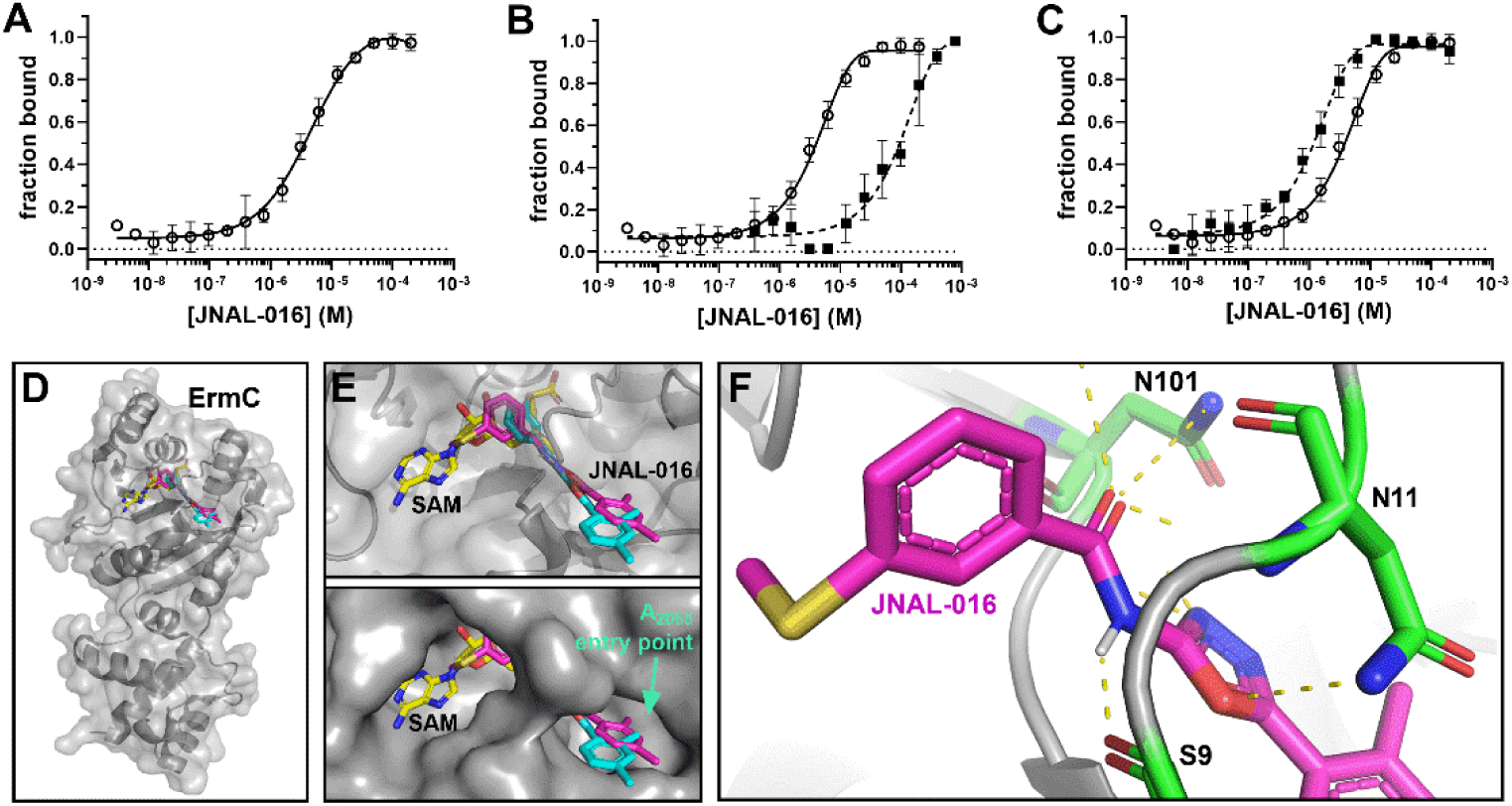
**JNAL-016 competes with SAM for the cofactor binding site**. Results from in vitro experiments highlighting the binding interaction between aza-SAM and JNAL-016 were determined using microscale thermophoresis. **A**) The binding profile of JNAL-016 (K_d_ = 1.8 ± 0.7 μM) to purified *Sa*ErmC (100 nM). **B**) Results from binding experiments to *Sa*ErmC showing the profile of JNAL-016 alone (open circles) and in the presence of 2 μM aza-SAM (K_d_ = 122 ± 13 μM; filled squares). **C**) Binding profiles for JNAL-016 alone (open circles) and in the presence of 45 μM 32-nt RNA substrate (K_d_ = 0.9 ± 0.2 μM; filled squares). Each data set represents averages of three replicates, and the error bars represent the standard deviation of the mean. **D**) Cartoon-surface representation of ErmC showing the docking results of JNAL-016 overlaid with a structure containing bound SAM within the cofactor binding pocket. The structure shows two different predicted conformations of the inhibitor, highlighted in cyan and magenta. **E**) Close-up view of the cofactor binding site showing the distinct predicted binding orientation for JNAL-016 relative to SAM. The top image shows a transparent surface view revealing the predicted conformations of the enzyme’s structure. The bottom image depicts the location of the binding pocket with a section of the molecule blocking the entry point for nucleotide A_2058_ during the methylation event, suggesting a possible inhibition strategy for the compound. **F**) Close-up showing a cartoon structure of ErmC docked with JNAL-016, highlighting the predicted polar contacts (yellow dashed lines) with specific residues of the enzyme, including N101, which is essential for substrate methylation during catalysis.

To confirm that JNAL-016 did not inhibit *Sa*ErmC catalysis competitively, we conducted a similar experiment, adding 45 μM of the 32-nt RNA substrate instead of aza-SAM. Unlike the effect observed with the cofactor, the presence of the RNA appeared to stabilize JNAL-016 binding, resulting in higher affinity (*_app_*K_d_ = 0.9 ± 0.2 μM) (**Figure 7C**). This observation implied that this compound was not competing with substrate binding, further supporting our enzyme inhibition kinetic data (**Figure 6C**). It is noteworthy that the latter observation could result from the large binding surface area of the substrate and the multiple enzyme-RNA contact points,^53^ which may hinder a competitive binding event with JNAL-016. Regardless, the presence of the RNA substrate resulted in a ∼2-fold enhancement in the affinity of JNAL-016 to *Sa*ErmC. In silico molecular docking experiments of this inhibitor to an AlphaFold modeled *Sa*ErmC structure localized it to the SAM binding site, with a section of the molecule potentially blocking the entry point of the adenine (A_2058_) to be methylated by the enzyme (**Figure 7D-F**). While a blockade at this location might result in competitive inhibition for a typical enzyme substrate, we propose that the large enzyme-substrate contact surface area would allow the substrate to remain tightly bound without being able to dock the nucleotide into the active site (**Figure S1 & S9**). Also, binding to the enzyme in the predicted conformation (**Figure 7E**) might explain the observed competition with SAM for the cofactor binding region. The predicted binding location for JNAL-016 has several implications, including its potential ability to act as a broad-spectrum adjuvant against ErmC variants. This proposal is based on the observation that currently identified ErmC variants have a significantly high sequence and structural conservation,^54^ especially when considering residues that make polar contacts with SAM within the cofactor binding pocket, including those that are essential for enzyme function, like N101, or ones that facilitate substrate binding (**Figure S9**).^54^ Taken together, these results further support our kinetic inhibition data demonstrating that JNAL-016 inhibits the activity of *Sa*ErmC noncompetitively. These findings also make a strong case for developing and using adjuvants as broad-spectrum inhibitors for tackling enzyme-mediated antibiotic resistance by ErmC variants.

## CONCLUSION

In this study, we investigated the use of small molecules to neutralize the activity of a variant of ErmC, which mediates resistance to MLS_B_ antibiotics in several pathogenic strains of bacteria. We successfully employed computer-aided solvent mapping and virtual screens to identify small-molecule binding sites and potential inhibitors of *Sa*ErmC. Using a developed *E. coli Sa*ErmC resistance model, we identified compounds with promising antibiotic-adjuvant properties against this form of resistance. Among these inhibitors, JNAL-016 displayed exceptional adjuvant activity when combined with ERY, enabling this antibiotic to regain its bactericidal properties, killing >99.9% of viable bacteria after 8 h. Our data characterizing this compound suggests it binds tightly to *Sa*ErmC, competing with SAM for the cofactor binding site to inhibit this enzyme noncompetitively. This comprehensive study offers a compelling case for prioritizing similar inhibitors to target resistance-conferring methyltransferases using antibiotic-adjuvant-based combination therapy to tackle deadly bacterial pathogens like *S. aureus*. The rapid spread of resistance-conferring methyltransferases like Erms in pathogenic strains of bacteria poses a significant threat to our ability to treat such infections.^31^ Given that the activity of such enzymes confers devastating resistance to clinically active antibiotic classes, it is essential to devise overarching countermeasures to deal with such cases. While this study may offer hope of a resolution, several key questions remain unanswered, warranting further investigation, including the potential clinical applications of Erm inhibitors or whether inhibitors like JNAL-016 could have broad-spectrum activity against other variants of this class of methyltransferases. Ongoing research will investigate how such compounds, and additional scaffolds, interact with these methyltransferases, providing vital insight into the design of more effective inhibitors with superior adjuvant properties.

## METHODS

### Computational solvent mapping

Solvent mapping and identifying small-molecule binding sites were conducted remotely using the FTMap servers (Boston University; http://ftmap.bu.edu/) as previously described^55^ with minor modifications. A PDB file of the available crystal structure of ErmC (PDB ID: 2ERC) without the cofactor bound was uploaded into the mapping algorithm to scan the entire structure. Probe-cluster consensus sites containing structurally diverse probes were used to identify predicted binding sites for small molecules.

### In silico virtual screening and molecular docking

Computational virtual screens were conducted using the AutoDock Vina computational software.^42^ The virtual antibacterial screening library containing 10,880 compounds was obtained from the Life Chemicals, Inc. repository as a compressed structural data file. This file was loaded onto the Open Babel software, which was used to conduct energy minimizations and transform the files into the required pdbqt file format for the docking algorithm. Similarly, the crystal structure of ErmC (PDB ID: 2ERC) was obtained from the Protein Data Bank and converted into a similar format using the AutoDock software. The docking grid search space was adjusted to cover the entire protein structure before initiating the docking event. The binding energies were used to rank and select the compounds used in this study, with prioritized compounds having the lowest energies and suitable drug-like properties projected using the Swiss ADME (absorption, diffusion, metabolism, and excretion) computational software.^43^

Molecular docking with JNAL-016 onto ErmC (PDB ID: 2ERC) was conducted using the AutoDock Vina software as previously described, with some modifications.^55^ The docking grid search space was adjusted to cover the entire protein structure before initiating the docking event. The optimal conformation(s) with the lowest binding energy and RMSD values = 0, were presumed to be the projected binding site for JNAL-016.

### Construction of the *E. coli Sa*ErmC resistance model and expression strains

The gene sequence of the *S. aureus ermC* (Uniprot ID: P02979) cloned into the pUC18 vector was constructed and obtained from Gene Universal Inc. Polymerase chain reaction was then used to amplify the gene with primers (forward primer 5’-ATA TAT GAA TTC ATG AAT GAA AAG AAC ATT AAA CAC; reverse primer: 5’-ATA TAT TCT AGA CTA TTT ATT GAA CAG TTT GTA CGA AT) designed to introduce restriction sites for cloning into the pBAD24 vector (5’-EcoRI/3’-Xbal). A 1% agarose gel was used to resolve the PCR product, and the desired band (∼ 756 bp) was excised and extracted from the gel fragments using the Monarch gel extraction kit (NEB). Double digests of the gene and cloning vector were performed using EcoRI and Xbal restriction enzymes in 50 µl reactions for 30 min at 37°C. The samples were resolved on a 1% agarose gel, and the desired fragments were isolated as highlighted above. T4 DNA ligase (NEB) was used to integrate *SaermC* into the digested vector, according to the manufacturer’s protocols, for 18 h at 16°C. The ligation product was transformed directly into chemical competent DH5α cells and plated on LB agar plates containing 100 μg/ml ampicillin (AMP). Plasmid was isolated from successful transformants, and the construct was confirmed by whole plasmid Sanger sequencing (Azenta), followed by transformation into chemically competent BW25113Δ*tolC::cat* cells.^56^ Successful transformants from this experiment resulted in strain TSB-001. The *Sa*ErmC purification strain, TSB-010, was constructed by transforming an IPTG-inducible pET28a plasmid containing an *S. aureus ermC* gene insert into BL21 (DE3) cells (constructed and obtained from Gene Universal).

### Broth microdilution assays to determine the minimum inhibitory concentrations (MICs) and half-maximal growth inhibitory concentrations (IC_50_s)

All broth microdilution screening assays followed the same procedures, with their respective antibiotics and/or inhibitory compounds. A 5 ml bacterial culture of strain TSB-001 or TSB-014 was prepared in LB containing 100 µg/mL AMP and grown overnight with shaking at 37°C. The next morning, the optical density at 600 nm (OD_600nm_) of a 10-fold dilution of the culture was determined. A bulk working culture was prepared by diluting the overnight culture to an OD_600nm_ of 0.004 (final OD_600nm_ = 0.002 in the assay). Next, 96-well plates were prepared with the bacterial culture containing antibiotics and/or test compounds. If *Sa*ErmC expression was required, 1% arabinose was added to the corresponding growth medium before assay set-up. Broth microdilution assays were performed as previously described with minor modifications.^46^ Typically, two-fold serial dilutions were performed in growth medium containing AMP in a 96-well plate, resulting in 50 μL of the test compound or antibiotic at 2x the desired final concentration in triplicate wells. Then, 50 μL of a culture at 2x the desired final cell density (OD_600nm_ = 0.002) was added to each well of the plates, which were then incubated at 37°C for 16-18 h. MIC values were determined by visual inspection of the plate to identify the lowest concentration without visible growth. To quantify the half-maximal inhibitory concentration (IC_50_) values, the OD_600nm_ of the samples was obtained using a SpectraMax^®^ iD5 plate reader (Molecular Devices). These data were analyzed relative to the ‘drug-free’ controls to generate normalized growth data. Triplicate data were averaged, and GraphPad Prism was used to generate dose-response growth inhibitory profiles, which were used to calculate the corresponding IC_50_ values for test compounds. At least three biological replicates were conducted for each experiment.

### Single-dose screening of ErmC inhibitory test compounds

A working stock culture of strain TSB-001 with 2x the desired final cell density was prepared as described above and supplemented with 1% arabinose. Three plates were prepared with LB broth: one with no antibiotic, one containing 1.25 µg/mL ERY, and one with 50 µg/mL CLN. The test compounds were added to triplicate wells for a final concentration of 0.02 µg/mL in the assay. The plates were incubated at 37 °C for 18 h, and the OD_600nm_ was recorded as described above. The raw data were normalized to the inhibitor-free controls (100% growth) and quantified in GraphPad Prism to generate a relative heat map for comparison.

### MIC determination using antibiotic strip diffusion assays

Solid top agar was liquified in a microwave, and 5 ml (containing 100 µg/mL AMP) was transferred into culture tubes and incubated at 44°C. 100 μL of a saturated overnight culture of strain TSB 001 was added to each sample along with 1% arabinose or 1% glucose (for expression or repression of ErmC, respectively). For experiments containing JNAL-016, it was added at 0.25 µg/mL into the molten top agar. The molten top agar samples were poured onto prewarmed LB-AMP plates and left at room temperature to solidify. Antibiotic test strips (Liofilchem) were then placed onto the solidified agar, and plates were incubated at 37°C for 20-24 h. MIC values were determined as the antibiotic concentration at which the growth clearing met the antibiotic strip. At least three biological replicates were obtained for each test, and the MICs were averaged.

### Checkerboard assays to assess the combined effects of erythromycin and JNAL-016

A 5 ml bacterial culture of strain TSB-001 or TSB-014 was prepared in LB containing 100 µg/mL AMP and grown overnight with shaking at 37°C. The next morning, the optical density (OD_600nm_) of a 10-fold dilution of the culture was determined. A bulk working culture was prepared by diluting the overnight culture to an OD_600nm_ of 0.004 (final 0.002). Next, a 96-well plate was prepared with LB supplemented with 100 µg/mL AMP and 2% arabinose for *Sa*ErmC expression (1% final in assay). ERY was serially diluted (2-fold) across the plate (columns 1-9, rows B-H), starting at 4x desired final concentration, and JNAL-016 down the plate (columns 2-9, rows A-H), beginning at 2x desired final concentration. Columns 10 and 11 were used to determine MICs of ERY and JNAL-016, respectively, on their own. Column 12 was used as a negative control in which DMSO was added instead of the antibiotic/test compound. Then, 50 μL of the culture at OD_600nm_ of 0.004 was added to each well of the plate, followed by incubation at 37°C for 18 h. MIC values were determined by visual inspection of the plate to identify the lowest concentration without visible growth. The fractional MICs were determined as the ratio of the MICs of ERY or JNAL-016 alone to the MIC of the combination. OD_600nm_ of the samples was obtained using a SpectraMax® iD5 plate reader (Molecular Devices), and heat maps were generated with GraphPad Prism. Three biological replicates were conducted, and the data were averaged to obtain the mean values. For strain TSB-001, the highest concentrations of ERY, CLN, and JNAL-016 used were 180 μg/mL, 4,250 μg/mL, and 1.0 μg/mL, respectively. Those for strain TSB-014 were 4,000 μg/mL, 5,312.5 μg/mL, and 50 μg/mL, respectively.

### Time-dependent growth, time-kill assays, and determination of the colony-forming units

For drug dilutions, a desired 2x concentration of JNAL-016 (final 0.085 µg/ml) was added to LB broth supplemented with 100 µg/mL AMP and 2% arabinose. Aliquots (96 µl) of this drug/antibiotic-containing medium were transferred to row A of a 96-well plate, and the other wells were filled with 50 µl of this sample. Then, 4 µl DMSO was added to wells A1 and A12, and 4 µl ERY (4 mg/ml) to wells A2-10. 2-fold serial dilutions were performed from row A through H, and 50 µL was discarded after the last well. A 2x bulk working culture (OD_600nm_ = 0.1) of strain TSB-001 supplemented with 100 µg/mL AMP and 10 µM PI fluorescent dye (dead cell staining) was prepared as described above. Next, 50 µL of this culture was added to all the wells, and the plate was incubated at 37 °C for 3 minutes with shaking (325 rpm). The T = 0 timepoint was obtained by recording the OD_600nm_ and PI fluorescence (excitation at 535 nm and emission at 615 nm) using the SpectraMax^®^ iD5 plate reader (Molecular Devices). The plate was then incubated at 37°C with shaking at 325 rpm for 8 h while monitoring the cell density and fluorescence readings every hour. The time-dependent growth curves and time-kill profiles were analyzed using GraphPad Prism. For the latter, the fluorescence values were normalized by the cell density of the corresponding sample-well before quantification. Determination of the colony-forming units (CFU/mL) was performed with the same samples by extracting aliquots (5 µL) at 0, 4, and 8-hour time points. The extracted volumes were diluted as follows, depending on the cell density recorded in the culture wells: cultures with an OD_600nm_ ≤ 0.15 were diluted 10^4^-fold, while those with a density > 0.15 were diluted 10^5^-fold using sterile water. Aliquots (50 µL) from these dilutions were then plated onto LB-AMP. The plates were incubated at 37°C for 24 h before colony counts. The data was analyzed using GraphPad Prism.

### Determination of the frequency of resistance

A 5 ml bacterial culture of strain TSB-001 was prepared in LB containing 100 µg/mL AMP and grown overnight with shaking at 37°C. Next morning, the culture was diluted by transferring 50 µL of the overnight culture into 5 mL of fresh LB media (100 µg/mL AMP) with 1% arabinose and 1 µg/mL ERY (for *Sa*ErmC expression). This culture grew until OD_600nm_ reached ∼ 0.8, then it was diluted to OD_600nm_ = 0.05 in LB. These cells were then diluted 10^1^, 10^2^, and 10^4^-fold in sterile nanopure water. Then, 50 µL aliquots from these diluted cells were spread onto warm LB plates. All plates were supplemented with 100 µg/mL AMP and 1% arabinose. The (+) ERY plates additionally had 50 µg/mL ERY. The (+) ERY and (+) JNAL-016 plates additionally had 50 µg/mL ERY and 0.042 µg/mL JNAL-016. The plates were incubated at 37°C for 24 h. Colonies were counted and converted to CFU/mL values. Frequency of resistance was calculated by dividing the CFU/mL values by the CFU/mL of the control plate. If no colonies were observed on the antibiotic selection plates, the spontaneous mutation frequency was determined as 1/total number of cells inoculated, indicating that the mutation frequency was lower than the detectable limit (1 CFU) under the experimental conditions. Three biological replicates were performed, and the averages (±SD) are reported.

### Cytotoxicity assays using the Human Embryonic Kidney cells [HEK-293]

The cell line was obtained from ATCC (CRL-1573). The cells were cultured in Eagle’s Minimum Essential Medium (EMEM) supplemented with 10% fetal bovine serum. Sterile black-clear bottom 96-well plates (Corning) were seeded with 1 x10^5^ cells/well and incubated at 37°C in a 5% CO_2_ incubator for 24 hours to allow the cells to adhere to the surface. After attachment, the media was aspirated off and replaced with the growth medium containing the different test compounds at concentrations equivalent to 20x their recorded MICs. Some wells received growth medium only, while others received media containing 1% DMSO as the positive control. The plates were incubated for 24 hours, as highlighted above, after which 10 μL of a lysis reagent (10mM Tris-Cl (pH 7.4), 150 mM NaCl, 1 mM EDTA, 1% Triton X-100) was added to select media-only wells. The plate was incubated in the 5% CO2 incubator for an additional 10 minutes before adding a solution of propidium iodide (5 μM final) in sterile water. The plate was incubated for an additional 20 min at 37°C in the 5% CO_2_ incubator before fluorescence measurements were recorded (excitation: 535 nm, emission: 615 nm) on a SpectraMax iD5 (Molecular Devices). The data were analyzed using GraphPad Prism and normalized to the lysis control (considered 100% lysed cells) to quantify the relative toxicity.

### Expression of recombinant *Sa*ErmC

ErmC was expressed by growing the TSB-010 strain first in 5 mL overnight cultures of LB containing 50 µg/mL kanamycin (KAN). Seed cultures were initiated by adding the overnight culture into 25 mL of Terrific Broth (TB) containing 50 µg/mL KAN and grown at 37°C shaking at 250 rpm for 3 h. The cultures were then added to 775 mL TB containing 50 µg/mL KAN and 1 µg/mL ERY (to ensure induction of ErmC), and grown at 37°C, 170 rpm to an OD_600nm_ ∼ 0.8. An uninduced sample was extracted, and expression of ErmC was induced with 1 mM IPTG for 16 h at 18°C, shaking at 150 rpm. An aliquot (1 mL) was extracted before harvesting. Cells were harvested by centrifugation at 4°C and 12,390 x g. The cell pellets were flash-frozen and stored at −80°C until use.

### Purification of recombinant *Sa*ErmC

The cell pellets were thawed on ice and resuspended in lysis buffer (50 mM Hepes, 500 mM NaCl, pH 7.5, 10% glycerol, 20 mM imidazole, 1 mM PMSF, 0.1 mg/mL lysozyme, 5 mM β-mercaptoethanol (β-ME)). The resuspension was stirred on ice for 30 minutes, then lysed by sonication. Cells were clarified by centrifugation at 39,200 x g (4°C). The supernatant was passed through an immobilized metal affinity chromatography (IMAC) nickel-nitrilotriacetic acid (Ni-NTA) column equilibrated with binding buffer (50 mM Hepes, 500 mM NaCl, pH 7.5, 10% glycerol, 20 mM imidazole, 5 mM β-ME). The column was washed with 5 column volumes of the binding buffer, and ErmC was eluted using an imidazole gradient ranging from 100, 200, 300, and 500 mM in the elution buffer (50 mM Hepes, 500 mM NaCl, pH 7.5, 10% glycerol, 5 mM β-ME). Elution fractions containing ErmC (assessed by SDS PAGE) were pooled together, concentrated, and buffer exchanged into the storage buffer (50 mM Hepes, 300 mM NaCl, pH 7.5, 10% glycerol, 5 mM β-ME) using a PD10 desalting column (Cytiva). *Sa*ErmC was purified further using a HiPrep 26/60 Sephacryl S-200 high-resolution size-exclusion column on the ÄKTA pure™ protein purification system (Cytiva), eluting with the storage buffer. Pooled fractions containing pure *Sa*ErmC were concentrated, and the concentration was quantified using the Bradford assay with bovine serum albumin (BSA) standards and estimated to be 3.8 ± 0.38 mg/mL (128 ± 12.8 µM). Aliquots of the pure protein were stored at −20°C.

### Generation of the SAH standard curve for the MTase-Glo assay

The MTase-Glo assay (Promega) was used to generate the calibration curve according to the manufacturer’s protocols. Briefly, 75 µL of a diluted sample of 1X reaction buffer (20 mM tris(hydroxymethyl)aminomethane (TRIS), pH 8, 50 mM NaCl, 1 mM ethylenediaminetetraacetic Acid (EDTA), 3 mM MgCl_2_, 0.1 mg/mL BSA, 1 mM dithiothreitol (DTT)) was added to each well in rows A-C, columns 2-12 of a 96-well sample preparation plate. 150 µL of 10 µM SAH was added to rows A-C, column 1. Two-fold serial dilutions were performed across the plate by transferring 75 µL from column 1 through column 11 and discarding excess sample (column 12 was left as a negative control with no SAH added). Aliquots (20 µL) from each serially diluted SAH concentration were transferred to the corresponding wells of an opaque, white flat-bottom 96-well assay plate. The plate was centrifuged for 1 min at 658 x g (RT). The MTase-Glo 10X reagent was diluted to 5X (according to the manufacturer’s directions), and 5 µL were added to all wells of the new plate (1X final in assay). The plate was centrifuged as described, shaken/mixed for 1 min, then incubated at RT for 30 mins. An aliquot (25 µL) of the MTase-Glo detection solution was added to all the wells (0.5X final in assay), followed by centrifugation as described above. The plate contents were mixed and incubated at RT for 30 min. Luminescence was measured at all wavelengths on the SpectraMax^®^ iD5 plate reader (Molecular Devices). The background luminescence from column 12 (containing no SAH) was subtracted from all luminescence measurements before data quantification. The SAH standard curve was generated with GraphPad Prism and was fit using linear regression (y = 169.3x – 809.8, R^2^ = 0.993). This same procedure was repeated with each new experiment to ensure an accurate SAH standard curve.

### Characterization of the *Sa*ErmC activity using the MTase-Glo assay

ErmC methylation activity was characterized using a 32-nt RNA oligomer substrate that mimics the part of the bacterial ribosome containing the A_2058_ target nucleotide for ErmC.^48^ This substrate was refolded before each experiment by denaturing at 65 °C for 10 minutes, then allowing the RNA to cool to RT slowly. The samples were prepared in 1X reaction buffer (see MTase-Glo assay description) using 100 µM 32-mer rRNA, 10 µM SAM, and 10 µM ErmC, with some sample tubes missing a single reaction component. They were incubated at RT, quenched after 1 h with 0.02 M HCl, and transferred to triplicate wells of an opaque, white flat-bottom 96-well assay plate. The MTase-Glo reagent and detection solution were added as described above to quantify the amount of SAM converted to SAH. The data was normalized to the sample containing all the reagents, which was set at 100% enzyme activity for the incubation period. Data from triplicate reactions were averaged to generate bar graphs using GraphPad Prism.

### Determination of the Michaelis-Menten kinetics for *Sa*ErmC catalysis (±) JNAL-016

All kinetics assays were performed at room temperature (RT), pH = 8, and atmospheric pressure. A 96-well plate was prepared with 1X reaction buffer (see above), 5 µM SAM, 32-mer rRNA concentrations ranging from 0.1 µM to 8 µM, either 100 µM JNAL-016 or 2% DMSO (control), and 0.5 µM *Sa*ErmC. Controls with no ErmC were prepared alongside the reactions to account for auto-hydrolysis of SAM. The reactions were quenched at: T = 1, 3, 5, 7, 10, 15, 30, 60, 90, 120 min by transferring 30 µL of reaction mix to wells with 7.5 µL of 0.1 M HCl at each time point (0.02 M HCl final). The MTase-Glo assay was used to quantify enzyme activity. The reactions were transferred to an opaque, white flat-bottom 96-well assay plate. The MTase-Glo Reagent was added at 1X final to all reaction wells and centrifuged for 1 min at 658 x g (RT). The plate was mixed for 1 min, then incubated at RT for 30 mins before adding the detection solution to all sample wells (0.5x final). The plate was centrifuged again, contents mixed and incubated at RT for 30 mins. The luminescence signal was measured and the data quantified as described above, using the SAH calibration curve. The kinetic assays were repeated 3 times, and the resulting data were quantified using GraphPad Prism.

### Binding affinity analyses using microscale thermophoresis

Purified ErmC was labeled via the C-terminal 6xHis-tag by incubating 200 nM of the protein with 200 nM NTA - Atto 647 N dye (Supelco) in phosphate-buffered saline, pH 7.4, with 0.05% Tween-20 (PBS-T) at room temperature (RT) for 30 mins, protected from light. The labeled ErmC was centrifuged at 15,000 x g, 4°C for 10 mins, and the supernatant was carefully transferred to a new microcentrifuge tube. Two-fold serial dilutions of JNAL-016 (from 400 µM to 0.012 µM) were prepared in PBS-T. For assays analyzing competitive binding of JNAL-016 with either aza-SAM (purchased from Antibodies.com) or 32-mer rRNA (re-folded as previously described), these latter reagents were added to the solution used to dilute the inhibitor, resulting in a final concentration of 2 μM or 45 µM, respectively. Labeled ErmC was added to each tube for a final concentration of 100 nM. Samples were loaded into microcapillary tubes, which were placed onto a sample holder and analyzed using the Monolith X (NanoTemper) to perform the binding affinity test with microscale thermophoresis. Binding experiments were performed at room temperature and repeated at least three times. The dissociation constants obtained from the replicates were averaged and reported with the corresponding standard deviation. GraphPad Prism was used to analyze the data and prepare the dose-dependent profiles with the “One-Phase Association” fit.

### Assessment of the expression levels of *Sa*ErmC using strain TSB-001

A 5 mL culture of strain TSB-001 and the parental strain (BW25113Δ*tolC::cat*) containing an empty pBAD24 vector was grown overnight at 37°C in LB-AMP. Next morning, the cultures were diluted to OD_600nm_ = 0.05 in LB-AMP medium supplemented with 1 µg/mL of ERY (to ensure induction of ErmC). The TSB-001 culture was split into 4 flasks (25 mL each). Three flasks were made with different percentages of arabinose to evaluate the expression levels of ErmC (0.2%, 0.5%, and 1%). The last flask was supplemented with 0.2% arabinose and 0.02 µg/mL JNAL-016 to determine if JNAL-016 impacts the expression level of ErmC. The cultures were incubated at 37°C, shaking at 250 rpm for 6 h. The cell density of each culture was determined, and they were all diluted to an OD_600nm_ = 1.0 before analysis using SDS PAGE.

### Sequence and structural alignments of the ErmC variants

The amino acid sequences of several ErmC variants were obtained from UniProt (IDs: P13956, P13978, P13957, P02979, Q79AA6, Q54284, Q4L2Y6, A0A2K0AWZ7, and A0A509LQ55). These sequences were aligned using the Muscle multiple sequence alignment (MSA) tool (EMBL-EBI) and WebLogo 3^57^ was used to generate a figure of the sequence conservation alignment. PyMoL was used to perform the structural alignments.

## SUPPORTING INFORMATION

The supporting information is available free of charge on the ACS Publications Website at DOI:

- Tables and figures highlighting additional results for computational experiments, cytotoxicity screens, antibacterial combinatorial assays, and protein expression, as well as materials used and compound identities

## Author contributions

J.N.A. conceived the study. J.N.A. performed cytotoxicity screens. T.S.B. and J.N.A. designed the experiments, analyzed the data, prepared, and wrote the manuscript.

## Notes

The authors declare no competing financial interest.

## ACKNOWLEDGEMENTS

We thank Annalee M. Schmidt for assistance in conducting the minimum inhibitory concentration assays for the initial 20 test compounds. This work was supported by start-up research funds to J.N.A. from the Department of Chemistry and Biochemistry at Miami University of Ohio.

## ABBREVIATIONS

SAM: *S*-adenosylmethionine
SAH: *S*-adenosylhomocysteine
Aza-SAM: 50-[N-[(3S)-3-Amino-carboxypropyl]-N-methylamino]-50-deoxyadenosine
Erm: Erythromycin resistance methyltransferase
ERY: Erythromycin
CLN: Clindamycin
TET: Tetracycline
PI: propidium iodide

