## Supplemental File for "Antibiotic-adjuvants abolish resistance conferred by the *Staphylococcus aureus* erythromycin resistance methyltransferase in an *Escherichia coli* model"

### SUPPORTING INFORMATION

#### Figure S1

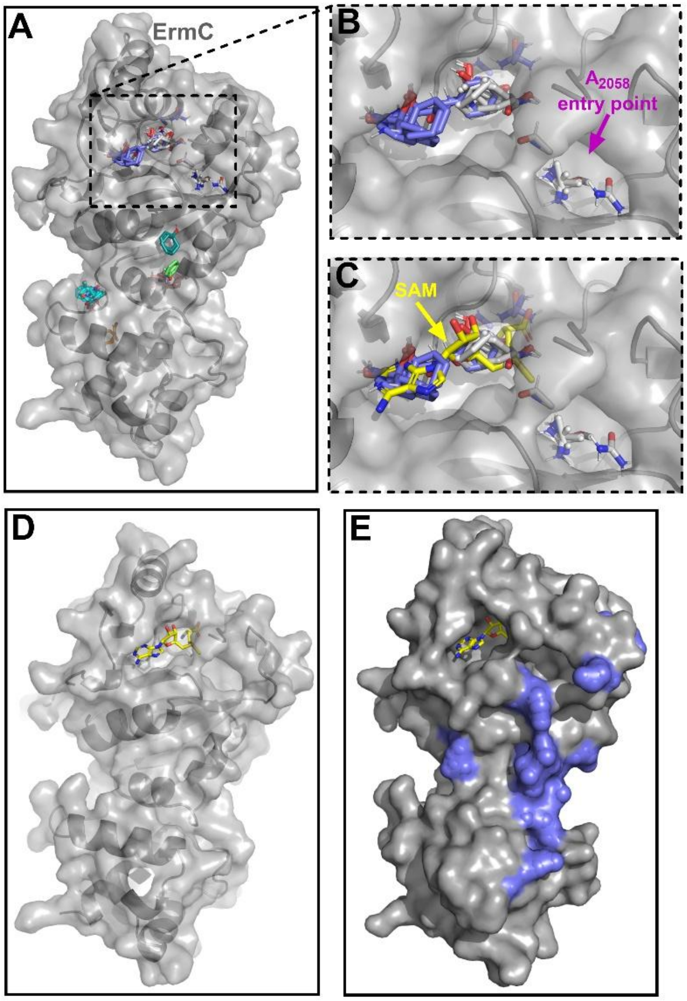


**Figure S1. Solvent mapping using FTMap accurately predicts the SAM binding site and identifies potential small-molecule binding hotspots**. **A**) Surface-cartoon structural representation of ErmC (PDB ID 2ERC) showing probe clusters (colored molecules) highlighting potential small molecule binding sites. **B**) Close-up view of the SAM binding and nucleotide ($A_{2058}$) entry points showing multi-structural probe cluster consensus sites (hotspots), validating the accuracy of this computational prediction method. **C**) Close-up view showing an overlaid structure of ErmC (PDB ID 1QAO) bound to SAM (yellow) and the AlphaFold predicted *Sa*ErmC model. The overlap of the cluster probes and SAM confirms its binding site. **D**) A surface-cartoon model showing the full structure of ErmC (PDB ID 1QAO) bound to SAM (yellow). **E**). Surface structure of the ErmC, highlighting the global distribution of the substrate RNA contact points.

### Figure S2

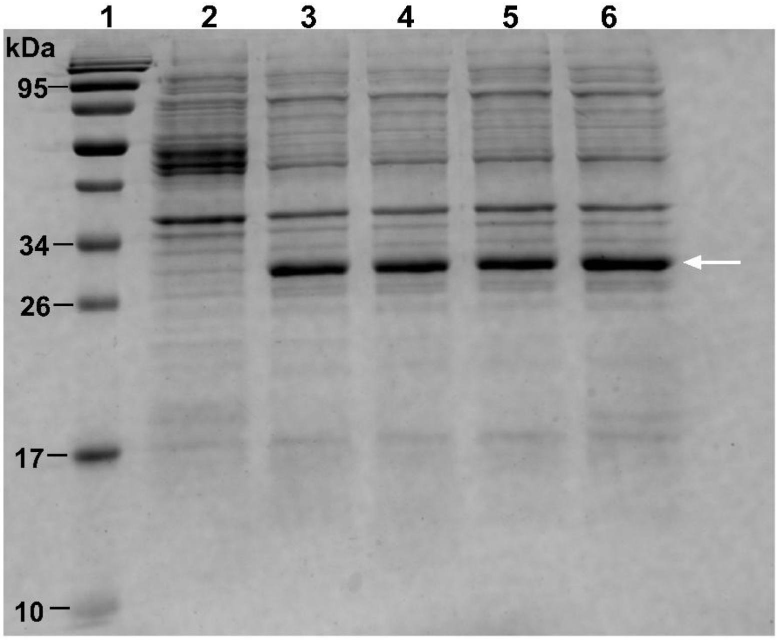


**Figure S2**. **SDS PAGE analysis gel confirms the expression of *Sa*ErmC in the *E. coli* resistance model strain upon induction with varying concentrations of arabinose (ARA)**. Lane ID: L1 – protein ladder; L2 – empty vector; L3 – 0.2% ARA; L4 – 0.5% ARA; L5 – 1% ARA; L6 – 0.2% ARA + 0.02 μg/ml JNAL-016. The band corresponding to the overexpressed *Sa*ErmC (~29 kDa) is shown with an arrowhead. Lanes 3 and 6 show similar expression levels in the presence or absence of JNAL-016.

### Figure S3

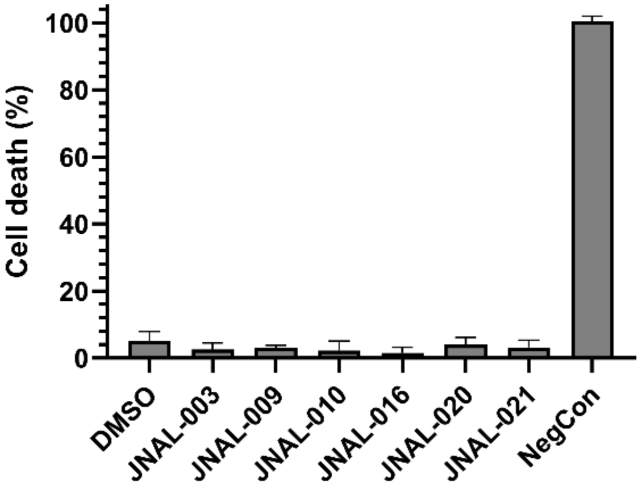


**Figure S3. Prospective adjuvants display no significant toxicity against HEK293 cells**. The test compounds were added to cells at 20x their MIC values (**Table S1**) and incubated for 24 h before analysis. The DMSO control represents the maximum volume (2%) of solvent added to the cultures in each sample well. The negative control (NegCon) contained 1x of the lysis reagent added after the 24 h incubation period. The data were normalized to the NegCon and are presented as the averages from three biological replicates ± the standard deviation.

### Figure S4

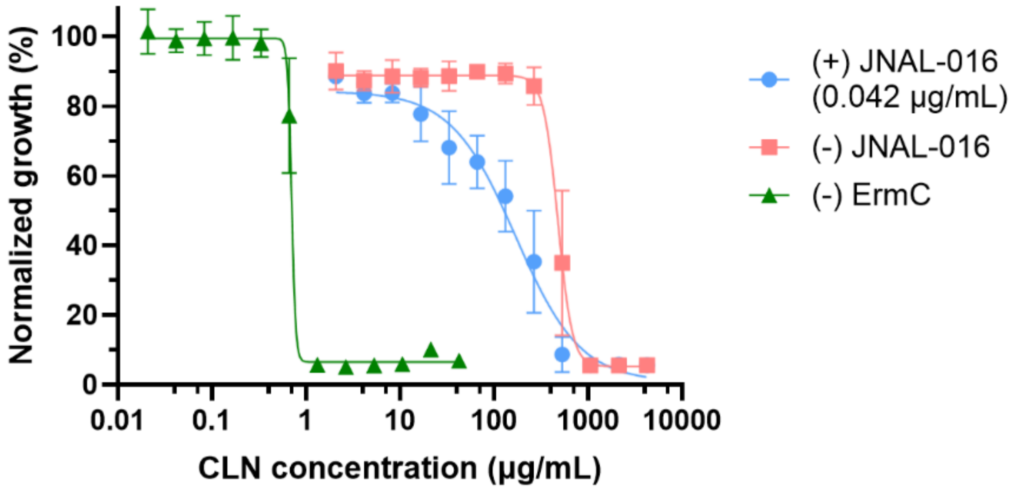


**Figure S4**. **JNAL-016 partially rescues the activity of CLN in the presence of *Sa*ErmC.** The graph shows the growth inhibitory dose response profiles for CLN assessed in the absence (green triangles) or presence (red squares) of *Sa*ErmC expression and in the presence of both *Sa*ErmC and JNAL-016 (0.042 μg/ml) (blue circles).

#### Figure S5

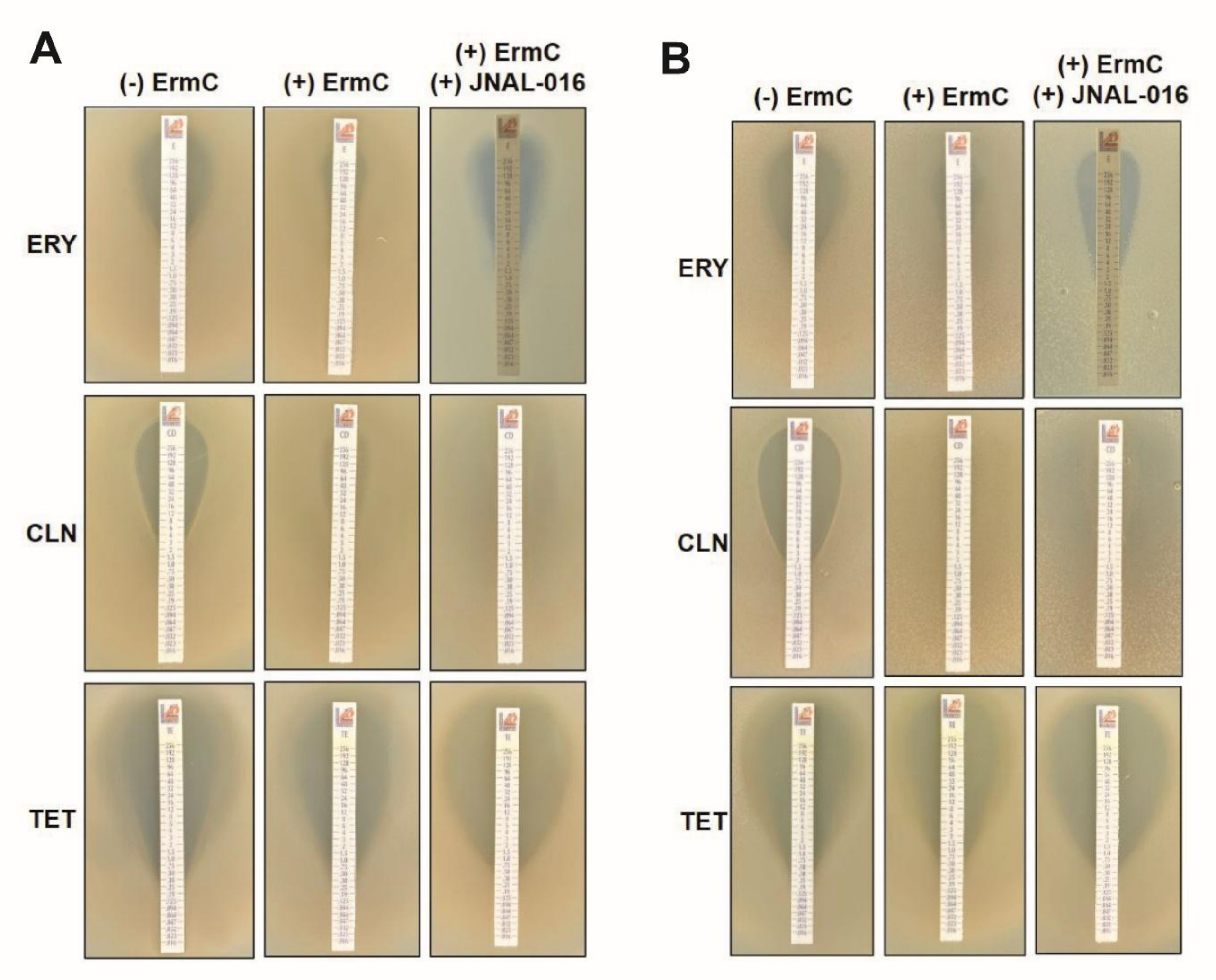


**Figure S5. Antibacterial activity determined using the top-agar method confirms the adjuvant properties of JNAL-016**. Biological replicates for the determination of MICs using antibiotic-infused strips. For induction of *Sa*ErmC and samples treated with the adjuvant, arabinose (1%) (middle column) or a combination of arabinose and JNAL-016 at 0.25 µg/ml (last column) were added to the molten top agar before solidifying it on the plate and adding the strips. The concentration of the compound (added to the top agar) used did not inhibit growth independently, based on the observed bacterial growth at the lower ends of the strips.

#### Figure S6

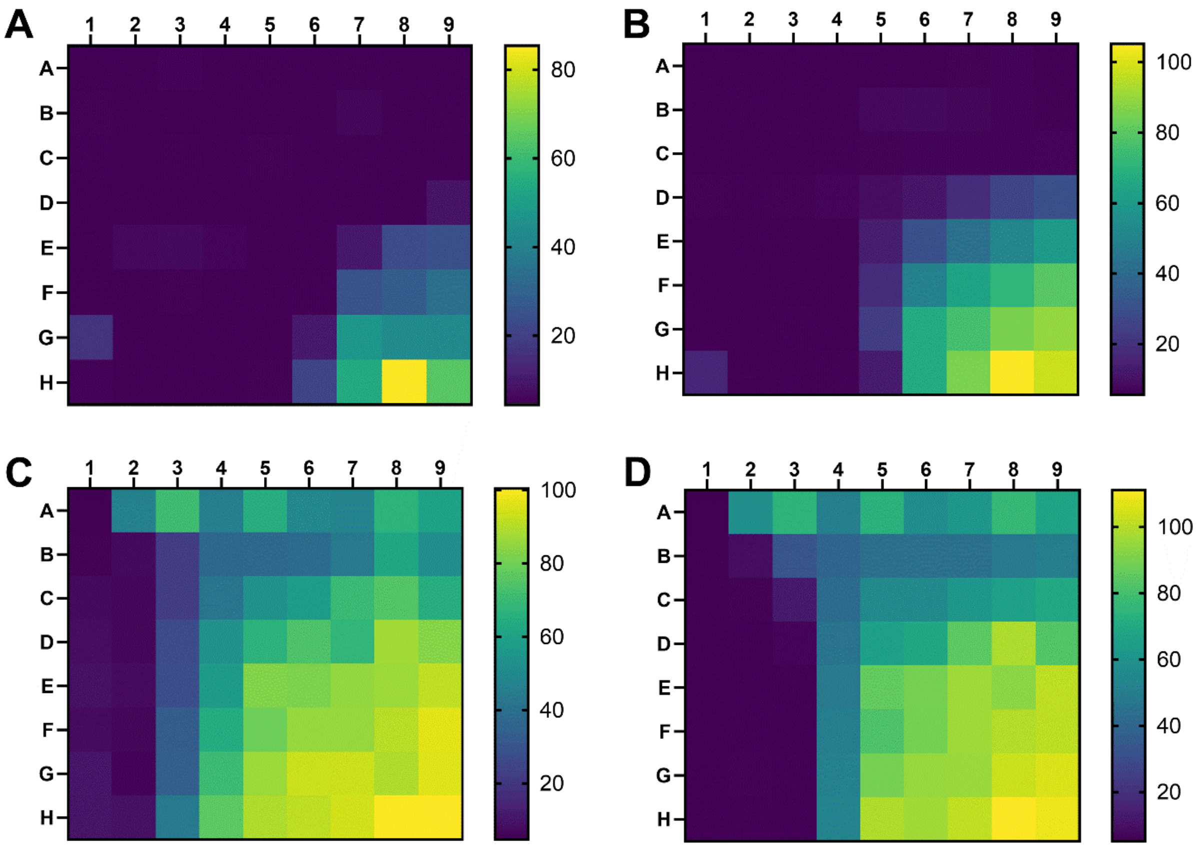


**Figure S6. Combinations of JNAL-016 with ERY or CLN act synergistically against ErmC-mediated resistance in the WT and Efflux-deficient strains**. Heat maps showing average results from checkerboard assays evaluating the growth inhibitory potential of combinations of JNAL-016 with ERY (**A** & **C**) or CLN (**B** & **D**) against the strain TSB-001 (TolC-deficient; **A** & **B**) or strain TSB-014 (WT; **C** & **D**) expressing *Sa*ErmC. ERY (or CLN) was serially diluted (2-fold) across the plate (columns 1-9, rows B-H), with the highest concentration at 2x MIC (final). JNAL-016 was serially diluted (2-fold) down the plates (columns 2-9, rows A- H), with the highest concentration at 2x MIC (final). The scale bar indicates normalized relative growth (%). The optical density data ($OD_{600nm}$) used to generate the heat map are averages from three independent experiments. The highest concentration of ERY in A = 180 μg/ml and C = 4,000 μg/ml. The highest concentration of CLN in B = 4,250 μg/ml and D = 5,312.5 μg/ml. Highest concentration of JNAL-016 in A & B = 1.0 μg/ml and C & D = 50 μg/ml.

### Figure S7

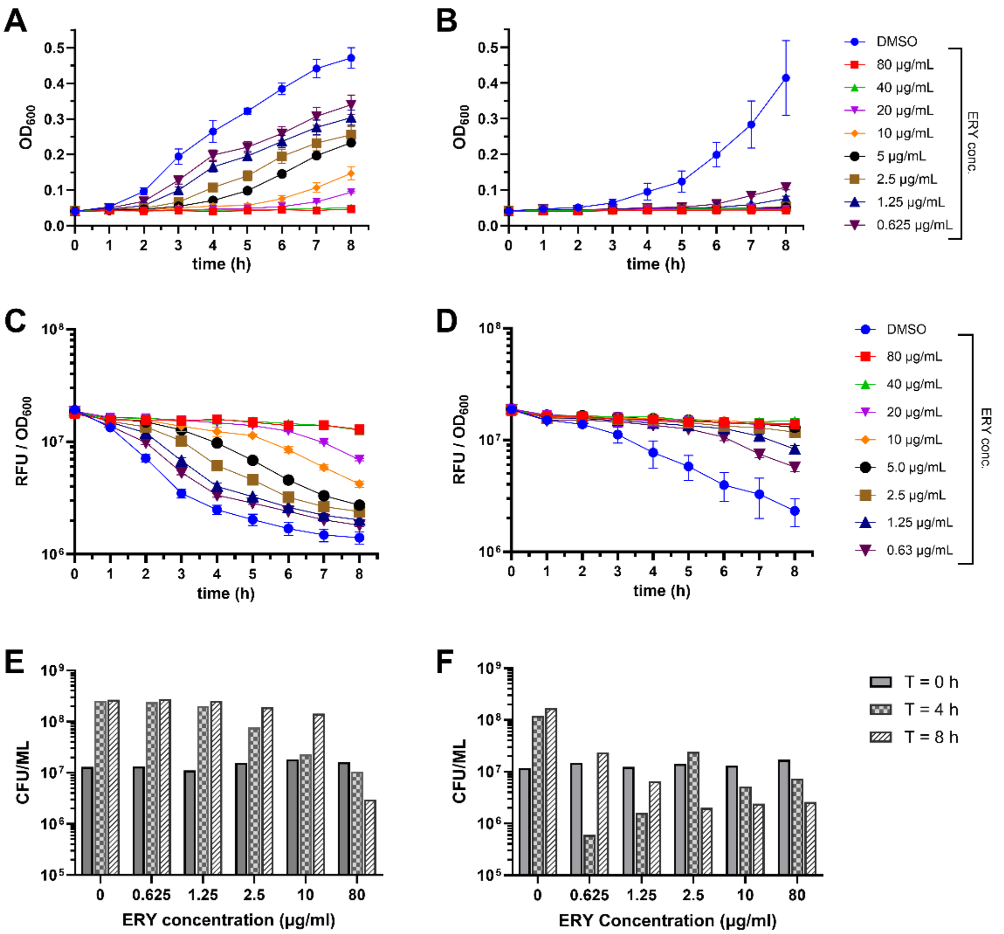


**Figure S7**. JNAL-016 acts as an adjuvant to restore the bactericidal properties of ERY in time-kill assays. **A & B**) Growth profiles of strain TSB-001 expressing *Sa*ErmC treated with varying concentrations of ERY monitored by cell density ($OD_{600\ nm}$) every hour over eight hours in the absence (**A**) or presence (**B**) of JNAL-016 (0.085 μg/ml) added to every sample. **C & D**) Time-kill profiles for the same samples in ‘A & B’ presented as cell-density-normalized PI fluorescence ($RFU/OD_{600\ nm}$) over eight hours in the absence (**C**) or presence (**D**) of JNAL-016. Samples with higher fluorescence on the graphs have more dead cells. **E & F**) Quantification of the colony-forming units from the same samples in ‘A & B’ performed at different timepoints and concentrations of ERY in the absence (**E**) or presence (**F**) of JNAL-016. DMSO control cultures for samples in panels A, C & E indicate no ERY present in the sample, while those in panels B, D, & F are treated with only JNAL-016 (dissolved in DMSO) at 0.085 μg/ml.

### Figure S8

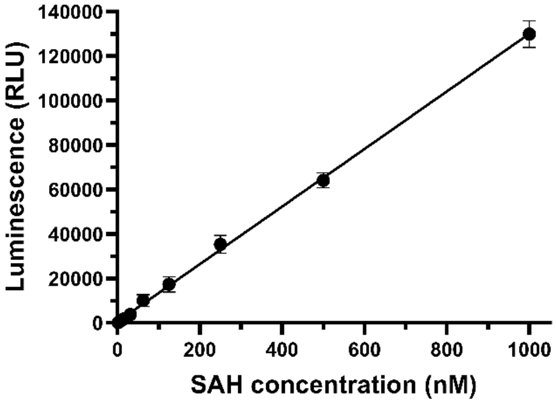


**Figure S8**. A representative calibration curve used to correlate the observed luminescence signal to the concentration of SAM transformed into SAH in the MTase-Glo *Sa*ErmC activity assays. This curve was regenerated alongside each new experiment to ensure accurate SAH concentration calculations.

#### Figure S9

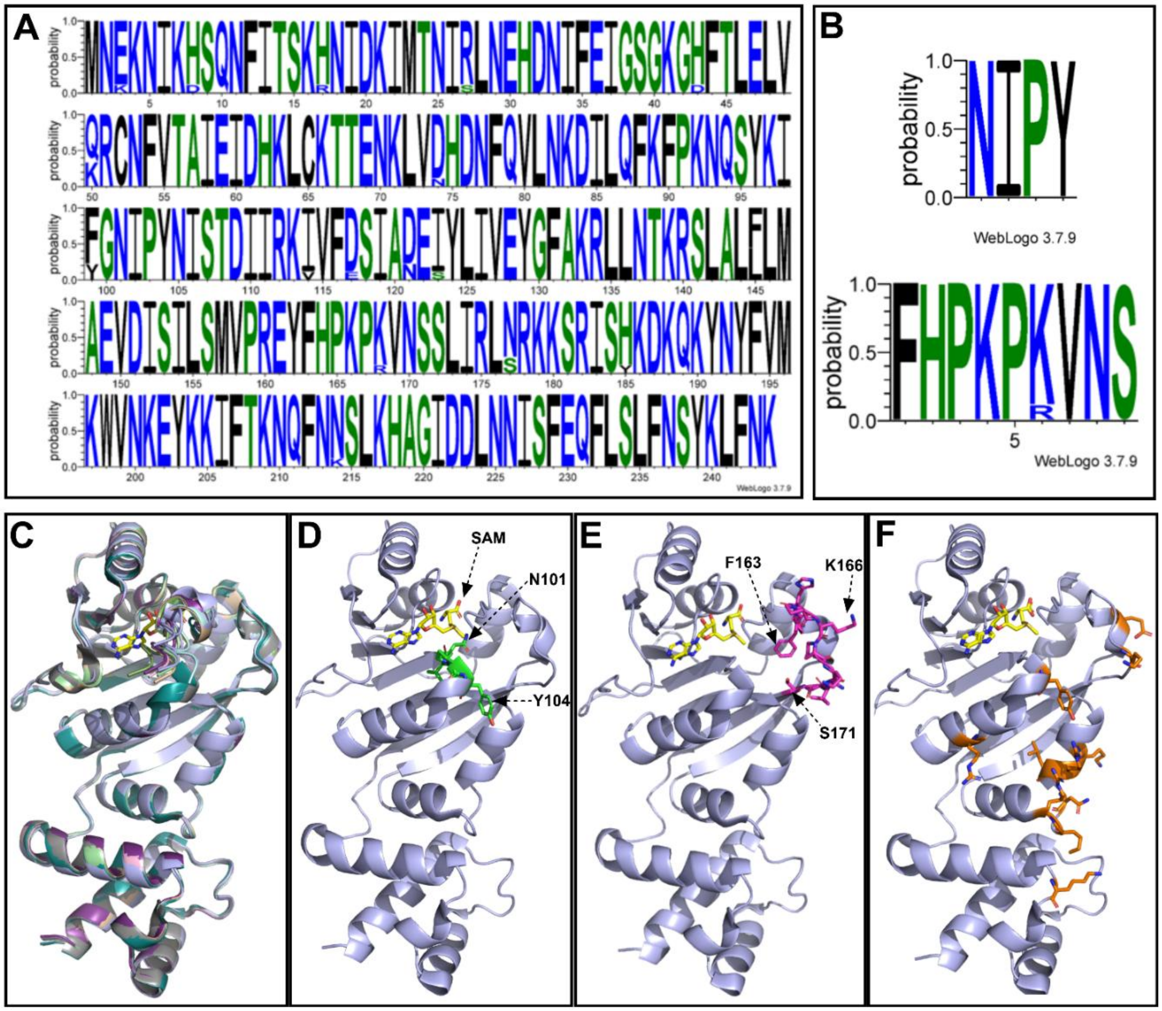


**Figure S9.** Multiple sequence and structural alignment highlighting the high amino acid conservation among ErmC variants. **A**) A WebLogo profile generated using ErmC variants from *B. subtilis* and several *S. aureus* species. UniProt IDs: P13956, P13957, P13978, P02979, Q79AA6, Q54284, Q4L2Y6, A0A2K0AWZ7, and A0A509LQ55. **B**) Segments of the WebLogo image highlighting sequence conservation of the four residues (N010, I102, P103, and Y104) essential for catalysis (top) and those making up the FXPXPXVXS motif demonstrated to be crucial for substrate modification by the enzyme (bottom). **C**) Cartoon representations of the structural alignment of documented ErmC variants used to generate the sequence comparison showing their high structural similarity. **D** & **E**) Cartoon representation of a representative ErmC structure (PDB ID: 1QAO), highlighting the location of some of the essential residues, NIPY (green, **D**) and FXPXPXVXS (magenta, **E**), required for catalysis. **F**). Cartoon representation of the ErmC structure (PDB ID: 1QAO) showing the locations of important residues (orange) that make up the large surface area of RNA substrate interaction points during catalysis.

### SUPPORTING INFORMATION TABLES

**Table S1.** Description of strains, plasmids, and cloning vectors used in this study.

| STRAIN | DESCRIPTION | SOURCE |
| --- | --- | --- |
| BW25113 (WT) | *E. coli* strain with *araBAD* deletion to inhibit arabinose metabolism. | ATCC[a] |
| BW25113Δ*tolC*::*cat* | BW25113 WT with *tolC* (efflux pump) deletion, replaced by the chloramphenicol resistance gene. | Main manuscript reference 55 |
| TSB-001 | BW25113Δ*tolC::cat* strain containing plasmid pBAD24-*ermC*. | This study |
| BL21(DE3) | *E. coli* protein expression strain. | NEB[b] |
| TSB-010 | BL21 (DE3) strain containing plasmid pET28a-*ermC*. | This study |
| TSB-014 | BW25113 (WT) containing plasmid pTSB-001 | This study |
| **PLASMID/VECTOR** | | |
| pBAD24 | Expression vector used for the controlled production of ErmC (arabinose - inducer/glucose - repressor). Contains an ampicillin resistance marker. | ATCC[a] |
| pET28a | Expression vector containing the *lac* operon for IPTG-inducible expression of a protein of interest. Used to attach a C-terminal 6xHis-tag to ErmC for purification. Contains a kanamycin resistance marker. | ATCC[a] |
| pTSB-001 | pBAD24 vector containing the *S. aureus ermC* gene for regulated expression of ErmC in cell-based assays. | This study |
| pTSB-002 | pET28a vector carrying the *S. aureus ermC* gene for the expression and purification of ErmC. | This study |

[a]ATCC – American Type Culture Collection
[b]NEB – New England Biolabs

**Table S2.** Recorded MICs for the 20 initial hits of prospective adjuvants used in the single-dose screens against strain TSB-001.

| Compound ID | MIC[a] (μg/ml) | | Compound ID | MIC[a] (μg/ml) |
|---|---|---|---|---|
| JNAL-001 | 0.9 (0.2) | | JNAL-012 | 0.3 (0.1) |
| JNAL-002 | >0.9 | | JNAL-013 | 0.7 (0) |
| JNAL-003 | >0.9 | | JNAL-014 | 0.7 (0.1) |
| JNAL-004 | >0.6 | | JNAL-015 | 0.7 (0) |
| JNAL-005 | 0.8 (0.2) | | JNAL-016 | 0.2 (0) |
| JNAL-006 | 0.7 (0) | | JNAL-017 | >0.84 |
| JNAL-008 | >0.7 | | JNAL-018 | 0.9 (0) |
| JNAL-009 | 0.2 (0) | | JNAL-019 | >1.0 |
| JNAL-010 | 0.1 (0) | | JNAL-020 | 0.2 (0) |
| JNAL-011 | 0.1 (0) | | JNAL-021 | 0.1 (0) |

[a]The data shown are averages from three or more independent experiments (SD).

**Table S3.** Information for the identities of the six test compounds used in this study was obtained from the antibacterial screening library (Life Chemicals, Inc.).

| COMPOUND ID | LIFE CHEMICALS, Inc. ID | CAS # | Smiles |
|---|---|---|---|
| **JNAL-003** | F0559-0394 | 533872-48-9 | CN(CC1=CC=CC=C1)S(=O)(=O)C1=CC=C(C=C1)C(=O)NC1=NN=C(O1)C1=CC=C(F)C=C1 |
| **JNAL-009** | F2518-0168 | 891144-28-8 | CC1=CC=C(C2=NN=C(NC(=O)C3=CC=C(C=C3)C(=O)C3=CC=CC=C3)O2)C(C)=C1 |
| **JNAL-010** | F2518-0186 | 891145-69-0 | CC1=CC=C(C2=NN=C(NC(=O)C3=CC=C(Cl)S3)O2)C(C)=C1 |
| **JNAL-016** | F2554-0165 | 898434-71-4 | CSC1=CC(=CC=C1)C(=O)NC1=NN=C(O1)C1=CC=C(C)C=C1C |
| **JNAL-020** | F2647-0161 | 898434-03-2 | CCSC1=CC=CC(=C1)C(=O)NC1=NN=C(O1)C1=CC=C(C)C=C1C |
| **JNAL-021** | F2647-0170 | 898459-29-5 | CCSC1=CC=CC(=C1)C(=O)NC1=NN=C(O1)C1=C(C)C=CC(C)=C1 |

**Table S4.** Antibacterial MIC and $IC_{50}$ profiles of CLN in the presence or absence of candidate adjuvants against strain TSB-001 expressing *Sa*ErmC.

| **Adjuvant Information** | | **CLN MIC[a] (μg/ml)** | | | **CLN $IC_{50}$[a] (μg/ml)** | | |
|---|---|---|---|---|---|---|---|
| **ID** | **Concentration used (μg/ml)** | **(-) adjuvant** | **(+) adjuvant** | **MIC Fold change** | **(-) adjuvant** | **(+) adjuvant** | **$IC_{50}$ Fold change** |
| **JNAL-003** | 0.058 | 1,062 ± 0 | 531 ± 0 | 2 | 522 ± 68 | 270 ± 37 | 1.9 |
| **JNAL-009** | 0.025 | | 531 ± 0 | 2 | | 233 ± 27 | 2.2 |
| **JNAL-010** | 0.010 | | 443 ± 153 | 2.4 | | 91 ± 74 | 5.7 |
| **JNAL-016** | 0.042 | | 443 ± 153 | 2.4 | | 165 ± 29 | 3.2 |
| **JNAL-020** | 0.022 | | 531 ± 0 | 2 | | 368 ± 114 | 1.4 |
| **JNAL-021** | 0.022 | | 531 ± 0 | 2 | | 271 ± 23 | 1.9 |

[a]The data shown are averages from three independent experiments ± SD.

**Table S5.** Antibacterial MIC values from antibiotic-infused strip assays assessed on solid LB agar plates against strain TSB-001.

| | MIC[a] (μg/ml) | | |
|---|---|---|---|
| **Antibiotic ID** | **(-) *Sa*ErmC** | **(+) *Sa*ErmC** | **(+) *Sa*ErmC/ (+) JNAL-016** |
| **ERY** | 7.3 ± 1.2 | 32 ± 2.4 | 3.0 ± 0.0 |
| **CLN** | 3.3 ± 1.2 | 256 ± 2.4 | 3.7 ± 0.6 |
| **TET** | 0.5 ± 0.0 | 0.67 ± 0.14 | 0.5 ± 2.4 |

[a]The data shown are averages from three independent experiments ± SD.

**Table S6.** Antibacterial MIC and $IC_{50}$ profiles of ERY and CLN in the presence or absence of JNAL-016 against strain TSB-014.

| | MIC[a] (μg/ml) | | | $IC_{50}$[a] (μg/ml) | |
|---|---|---|---|---|---|
| **Antibiotic** | **(-) ErmC** | **(+) ErmC** | **(+) ErmC, (+) 0.4 μg/ml JNAL-016** | **(-) ErmC** | **(+) ErmC** |
| ERY | 250 ± 0 | 8,000 ± 0 | 2,000 ± 0 | 57.8 ± 1.0 | 1,877 ± 362 |
| CLN | 93.8 ± 0 | 2,656 ± 0 | 1,328 ± 0 | 53.1 ± 4.5 | 922 ± 165 |

[a]The data shown are averages from three independent experiments ± SD.
